# EXP1 is required for organization of the intraerythrocytic malaria parasite vacuole

**DOI:** 10.1101/752634

**Authors:** Timothy Nessel, John M. Beck, Shima Rayatpisheh, Yasaman Jami-Alahmadi, James A. Wohlschlegel, Daniel E. Goldberg, Josh R. Beck

## Abstract

Intraerythrocytic malaria parasites reside within a parasitophorous vacuole membrane (PVM) that closely overlays the parasite plasma membrane (PPM) and constitutes the barrier between parasite and host compartments. The PVM is the site of several essential transport activities but the basis for organization of this membrane system is unknown. We utilized the second-generation promiscuous biotin ligase BioID2 fused to EXP2 or HSP101 to probe the content of the PVM, identifying known and novel candidate PVM proteins. Among the best represented hits were members of a group of single-pass integral membrane proteins that constitute a major component of the PVM proteome but whose function remains unclear. We investigated the function of EXP1, the longest known member of this group, by adapting a CRISPR/Cpf1 genome editing system to install the TetR-DOZI-aptamers system for conditional translational control. EXP1 knockdown was essential for intraerythrocytic development and accompanied by profound changes in vacuole ultrastructure, including increased separation of the PVM and PPM and formation of abnormal membrane structures in the enlarged vacuole lumen. While previous *in vitro* studies indicated EXP1 possesses glutathione S-transferase activity, a mutant version of EXP1 lacking a residue important for this activity *in vitro* still provides substantial rescue of endogenous *exp1* knockdown *in vivo*. Intriguingly, while activity of the *Plasmodium* translocon of exported proteins was not impacted by depletion of EXP1, the distribution of the translocon pore-forming protein EXP2 was substantially altered. Collectively, our results reveal a novel PVM defect that indicates a critical role for EXP1 in maintaining proper PVM organization.

**Importance:** Like other obligate intracellular apicomplexans, blood-stage malaria parasites reside within a membrane-bound compartment inside the erythrocyte known as the parasitophorous vacuole. Although the vacuole is the site of several transport activities essential to parasite survival, little is known about its organization. To explore vacuole biology, we adopted recently developed proteomic (BioID2) and genetic (CRISPR/Cpf1) tools for use in *Plasmodium falciparum*, which allowed us to query the function of the prototypical vacuole membrane protein EXP1.

Knockdown of EXP1 showed that a previously reported glutathione S-transferase activity cannot fully account for the essential function(s) of EXP1 and revealed a novel role for this protein in maintaining normal vacuole morphology and PVM protein arrangement. Our results provide new insight into vacuole organization and illustrate the power of BioID2 and Cpf1 (which utilizes a T-rich PAM uniquely suited to the *P. falciparum* genome) for proximity protein identification and genome editing in *P. falciparum*.

## Introduction

The obligate intracellular malaria parasite *Plasmodium falciparum* resides within a parasitophorous vacuole (PV) established during host cell invasion which constitutes the principal barrier between the parasite and its host cell (1). During the parasite blood stage, several essential transport activities at this membrane enable host erythrocyte subversion and parasite growth (2). This includes the uptake of large amounts of host cytosol through a process involving a double-membrane invagination of the parasitophorous vacuole membrane (PVM) and parasite plasma membrane (PPM) known as the cytostome (3). Catabolism of hemoglobin, the principal component of this ingested material, provides free amino acids for parasite metabolism and opens space within the host compartment for parasite expansion (4–6). The vacuole is also a key trafficking site for the export of hundreds of parasite proteins which are first secreted into the PV and then translocated across the PVM by the *Plasmodium* translocon of exported proteins (PTEX) (7–9). The PTEX core complex is composed of the HSP101 AAA+ ATPase chaperone, which unfolds exported cargo and passes it through an oligomeric pore in the PVM formed by a second PTEX protein, EXP2 (10). A third component, PTEX150, serves to couple HSP101 to the pore. While proteins must be unfolded to pass through the EXP2 pore (11), small molecules up to 1,400 Da can pass through this channel, allowing for nutrient uptake and waste exchange (12–14).

Aside from EXP2, the known *P. falciparum* PVM proteome is comprised mainly of a group of single-pass integral membrane proteins oriented with C-terminus facing the host cytosol and N-terminus in the PV lumen (15–17). This group consists of the prototypical PVM protein EXP1 and the early transcribed membrane proteins (ETRAMPs, also known as small exported proteins (SEPs) in *P. berghei*) and contains some of the most highly expressed genes in the blood stage (17–19). The function of these proteins is largely unknown although EXP1 has been shown to possess glutathione S-transferase (GST) *in vitro*, which has been proposed to support detoxification of hematin released by hemoglobin catabolism (20).

One of the most striking features of the PV is the intimate proximity of the PVM and PPM, which is maintained until a very late stage of parasite development when PVM rounding occurs just prior to egress (2, 21). Additional lateral organization of this compartment is suggested by the formation of distinct oligomeric arrays of EXP1 and ETRAMPs in the PVM (22) and by the non-uniform distribution of PTEX components as well as PV-targeted exported and non-exported fluorescent fusion reporter proteins, which have been shown to display a punctate distribution in the PV described as a “necklace of beads” (7, 23–27). Visualized by immunofluorescence, this punctate arrangement is most prominent in the early ring stage of parasite development and resolves into a more homogenous distribution in the trophozoite and schizont stage, particularly for EXP2 (26). While these observations point to a highly coordinated membrane system, the basis for PV/PVM organization is unknown.

Here, we applied the second generation BioID2 proximity ligase system fused to multiple components of PTEX to probe the protein content of the PV and PVM, revealing known vacuole proteins as well as novel PV/PVM candidates. To interrogate the function of select hits, we adapted a CRISPR/Cpf1 system that recognizes a T-rich protospacer adjustment motif, greatly expanding the repertoire of guide RNA targets available for editing the *P. falciparum* genome.

Prompted by the high representation of EXP1 in our BioID2 datasets, we used Cpf1 editing to generate an EXP1 conditional knockdown mutant to explore its function. Depletion of EXP1 produced a lethal defect that was largely rescued by a mutant version of EXP1 shown to be defective in GST activity i*n vitro*, suggesting that this activity does not fully account for EXP1 function *in vivo*. Instead, EXP1 knockdown resulted in dramatic changes to PV morphology and PVM protein organization, revealing a novel role in maintaining proper order within the PV.

## Materials and Methods

### Parasite Culture

Deidentified, IRB-exempt red blood cells (RBCs) were obtained from the American National Red Cross. *P. falciparum* NF54^attB^ and derivatives were maintained under 5% O_2_, 5% CO_2_, and 90% N_2_ at 2% hematocrit in RPMI 1640 supplemented with 27 mM sodium bicarbonate, 11 mM glucose, 0.37 mM hypoxanthine, 10 μg/ml gentamicin ad 0.5% Albumax I (Gibco).

### Plasmids and genetic modification of *P. falciparum*

Cloning was carried out with Infusion (Clontech) or NEBuilder HiFi (NEB) unless noted otherwise. Primer and synthetic gene sequences are given in Table S1. To generate a BioID2 fusion to the endogenous EXP2 C-terminus, the coding sequence of the *Aquifex aeolicus* biotin ligase with an R40G mutation (BioID2) bearing a 3’ 3xHA epitope tag was amplified with primers P1/P2 from plasmid MCS-BioID2-HA (28) (Addgene #74224) and inserted between AvrII and EagI in plasmid pyPM2GT-EXP2-mNeonGreen (29), replacing the AvrII site with an NheI site and resulting in the plasmid pyPM2GT-EXP2-BioID2-3xHA. This plasmid was linearized at the AflII site between the 3ʹ and 5ʹ homology flanks and co-transfected with pUF-Cas9-EXP2-CT-gRNA (14) into NF54^attB^. Selection was applied with 2 μM DSM1 (30) 24 hours post-transfection. A clonal line was isolated by limiting dilution after parasites returned from selection and designated NF54^attB^::EXP2-BioID2-3xHA.

For fusion of BioID2 to endogenous HSP101, a flank assembly targeting the 3’ end of *hsp101* was amplified from plasmid pPM2GT-HSP101-3xFlag (14) with primers P3/P4 and inserted between XhoI and NheI in pyPM2GT-EXP2-BioID2-3xHA, resulting in plasmid pyPM2GT-HSP101-BioID2-3xHA. This plasmid was linearized at AflII, co-transfected with plasmid pAIO-HSP101-CT-gRNA (14) into NF54^attB^, selected with 2 µM DSM1 and cloned upon returning from selection, resulting in the line NF54^attB^::HSP101-BioID2-3xHA.

For Cas9-mediated editing of the *exp1* locus, the pAIO (31) plasmid was first simplified by removing the yDHODH-2A fusion to Cas9-NLS-FLAG using a QuikChange Lightning Multi Site Directed Mutagenesis kit (Agilent) and the primer P5, resulting in the plasmid pAIO2. The BtgZI site in the pre-sgRNA cassette was then replaced with AflII by inserting the annealed sense and antisense oligo pair P6/P7 into BtgZI, resulting in the plasmid pAIO3. Three Cas9 targets were chosen at the 3’ end of *exp1* (GACGACAACAACCTCGTAAG, AGGTTGTTGTCGTCACCTTG and AGTGTTCAGTGCCACTTACG) and each guide RNA (gRNA) seed sequence was synthesized as a sense and anti-sense oligo pair (P8/P9, P10/P11, and P12/P13, respectively). Each oligo pair was annealed and inserted into the AflII site of pAIO3 to yield the plasmids pAIO3-EXP1-CT-gRNA1, pAIO3-EXP1-CT-gRNA2 and pAIO3-EXP1-CT-gRNA3.

For Cpf1 editing, AsCpf1 and LbCpf1 were PCR amplified from plasmids pcDNA3.1-hAsCfp1 and pcDNA3.1-hLbCfp1 (Addgene #69982 and #69988) (32) with primer pairs P14/15 and P16/15, respectively, and inserted into pAIO3 between BamHI and XhoI. The Cas9 pre-gRNA cassette was then replaced with a pre-gRNA cassette with the appropriate direct repeat region for AsCpf1 or LbCpf1 using a QuikChange Lightning Multi Site Directed Mutagenesis kit and the primers P17 or P18, resulting in plasmids pAIO-AsCpf1 and pAIO-LbCpf1, respectively. For Cpf1-mediated editing of the *exp1* locus, two Cpf1 targets were chosen at the 3’ end of *exp1* (CAGCTGTTTAGTGTTCAGTGCCAC and GTGTTCAGTGCCACTTACGAGGTT) and each gRNA seed sequence was synthesized as a sense and anti-sense oligo pair for AsCpf1 (P19/P20 and P21/P22, respectively) and LbCpf1 (P23/P24 and P25/P26, respectively). Each oligo pair was annealed and inserted into the AflII site of pAIO-AsCpf1 or pAIO-LbCpf1 to yield the plasmids pAIO-AsCpf1-EXP1-CT-gRNA1, pAIO-AsCpf1-EXP1-CT-gRNA2, pAIO-LbCpf1-EXP1-CT-gRNA1 and pAIO-LbCpf1-EXP1-CT-gRNA2. To generate selectable Cpf1 plasmids, the AflII site between the yDHODH and Cas9 expression cassettes in the plasmid pUF-Cas9-pre-sgRNA (14) was replaced with an XmaI site using QuikChange Lightning Multi Site Directed Mutagenesis kit and the primer P27. The Cpf1 coding sequence with the PbDT 3’ UTR and adjacent pre-gRNA cassette was then amplified from pAIO-AsCpf1 or pAIO-LbCpf1 using primers P28/P29 or P28/P30 and inserted between XhoI and NotI, resulting in the plasmids pUF-AsCpf1-pre-gRNA and pUF-LbCpf1-pre-gRNA, respectively. Oligo pairs P21/P22 and P25/P26 were annealed and inserted into the AflII site in the pre-gRNA cassette of these vectors to yield pUF-AsCpf1-EXP1-CT-gRNA2 and pUF-LbCpf1-EXP1-CT-gRNA2, respectively.

For fusion of 3xHA-GFP11 to EXP1, a 5’ homology flank (up to, but not including, the stop codon) was amplified from NF54^attB^ genomic DNA using the primers P31/P32. As the *exp1* coding sequence did not allow for synonymous mutations in the protospacer adjustment motifs of the three Cas9 gRNAs, several synonymous mutations were incorporated in the seed sequences of these target sites. A 3’ homology flank (beginning 11 bp downstream of the stop codon) was amplified using the primers P33/P34 and the flank amplicons were assembled in a second PCR reactions using the primers P32/P33 and inserted between XhoI and AvrII in pyPM2GT-EXP2-3xHA-GFP11 (14), resulting in the plasmid pyPM2GT-EXP1-3xHA-GFP11. This plasmid was linearized at the AflII site between the 3ʹ and 5ʹ homology flanks and co-transfected into NF54^attB^ with the above Cas9 and Cpf1 plasmids designed to target *exp1*. Selection was applied with DSM1 and the expected integration was confirmed by diagnostic PCR using primers P35/P36.

For generation of the ETRAMP10.2^apt^ and ETRAMP5^apt^ lines bearing a 3xHA fusion, a Cpf1 gRNA target was chosen just upstream of the *etramp10.2* and *etramp5* stop codons (TGACTCTTGGTGTGGTACTTCTTC and GGTTCTTCGGTTTTGACTTCGTCT, respectively) and the gRNA seed sequences were synthesized as the sense and anti-sense primer pairs P37/P38 and P39/P40, which were annealed and inserted into the AflII site of the plasmid pAIO-AsCpf1, resulting in the plasmids pAIO-AsCpf1-ETRAMP10.2-gRNA1 and pAIO-AsCpf1-ETRAMP5-gRNA1. A modified version of the plasmid pMG75 (33) was generated by first replacing the AvrII site with an NcoI site using a QuikChange Lightning Multi Site Directed Mutagenesis kit and the primer P41. The BirA*-3xHA coding sequence from plasmid pBirA*-3xHA-LIC-DHFR (34) was then amplified with primers P42/P43 and inserted between BstEII/AatII and the 2x*attP* sequence was subsequently removed by using QuikChange Lightning Multi Site Directed Mutagenesis kit and the primer P44, resulting in the plasmid pMG75ΔattP-BirA*-3xHA. To target the 3’ end of *etramp10.2* or *etramp5*, a 5’ homology flank (up to but not including the stop codon) was amplified from NF54^attB^ genomic DNA using primers pairs P45/P46 or P47/P48, incorporating synonymous mutations in the seed sequence and protospacer adjustment motif of the gRNA target sites within each *etramp* coding sequence. A 3’ homology flank (beginning 89 or 257 bp downstream of the stop codon, respectively) was amplified using the primer pairs P49/P50 or P51/P52. The corresponding 5’ and 3’ flank amplicons were assembled in a second PCR reaction using the primer pairs P46/P49 and P48/P51 and inserted between AscI/AvrII in pMG75ΔattP-BirA*-3xHA, removing the BirA* sequence and resulting in the plasmids pMG75ΔattP-ETRAMP10.2-3xHA and pMG75ΔattP-ETRAMP5-3xHA. These plasmids were linearized at the AflII site and co-transfected with the corresponding pAsCpf1-gRNA plasmid into NF54^attB^ parasites. Cultures were maintained with 1µM aTc from the time of transfection and selection with 2.5 µg/ml Blasticidin-S was applied 24 hours post-transfection. After returning from selection, parasites were cloned and proper integration at the 3’ end of *etramp10.2* or *etramp5* was confirmed by PCR with primer pairs P53/P36 or P54/P36, respectively.

For generation of a combined TetR-DOZI-aptamers (TDA) and DiCre knockdown system to target *exp1*, the tandem NLS-FKBP12-Cre19-59 and NLS-FRB-Cre60-343 cassettes from plasmid pDiCre (35) were amplified with primers P55/P56 and inserted into the AscI site of plasmid pSN054 (36), a pJAZZ-based plasmid containing the TDA elements. This plasmid also contains *loxP* sites immediately upstream of the 5’ aptamer and immediately downstream of the TetR-DOZI cassette. The latter *loxP* site is adjacent to the AscI site and was removing during insertion of the DiCre cassette. To target *exp1*, 5’ and 3’ homology flanks immediately upstream and downstream of the *exp1* start and stop codons were amplified with the primer pairs P57/P58 and P59/P60 and inserted sequentially at FseI (5’ flank) and between I-CeuI/I-SceI (3’ flank). A promoterless *mruby3* coding sequence with the *hsp86* 3’ UTR was amplified from plasmid pLN-HSP101-SP-mRuby3 (21) using primers P61/P62, adding a *loxP* site immediately before the *mruby3* start codon, and inserted at I-CeuI. Finally, the *exp1* coding sequence (without introns) was recoded, synthesized as a gene block (IDT) and PCR amplified with primers P63/P64, adding a 3’ 3xHA tag, and inserted at AsiSI, resulting in the plasmid pEXP1^apt^. This plasmid was co-transfected with pAIO-LbCpf1-EXP1-CT-gRNA1 into NF54^attB^::HSP101-3xFLAG (14). Parasites were maintained with 1µM aTc from the time of transfection and 2.5 µg/ml Blasticidin-S was applied 24 hours later. A clonal line was isolated by limiting dilution after parasites returned from selection and designated EXP1^apt^. Integration at the 5’ and 3’ ends of the *exp1* locus was evaluated by PCR with primer pairs P65/P36 and P66/P67, respectively. Excision by DiCre was monitored with the primers P65/P68.

For complementation of EXP1^apt^ parasites, the re-coded *exp1* coding sequence (without introns) was PCR amplified from the synthesized gene block with primers P69/P70 and inserted between XhoI and NheI in pyEOE-attP-EXP2-3xMYC (14), resulting in the plasmid pyEOE-attP-EXP1-WT-3xMYC. The *exp1* codon 70 in this plasmid was then changed from AGA to ACA using a QuikChange Lightning Multi Site Directed Mutagenesis kit and the primer P71, resulting in the plasmid pyEOE-attP-EXP1-R70T-3xMYC. Finally, the mNeonGreen coding sequence was amplified from pyPM2GT-EXP2-mNeonGreen (29) with primers P72/P73 and inserted at NheI in pyEOE-attP-EXP1-WT-3xMYC, resulting in the plasmid pyEOE-attP-EXP1-mNeonGreen-3xMYC. Complementing plasmids were co-transfected with pINT (37) into EXP1^apt^ parasites to facilitate integration into the *attB* site on chromosome 6 and selection with 2 µM DSM1 was applied 24 hours post-transfection (in addition to 2.5 µg/ml Blasticidin-S and 1µM aTc for maintenance of endogenous *exp1* control by the TetR-DOZI-aptamers system). Parasites were cloned when they returned from selection and expression of EXP1 second copies was confirmed by Western blot.

For targeting mRuby3 to the PV, the *exp2* promoter and signal peptide were amplified from NF54^attB^ genomic DNA with primers P74/P75 and inserted between AatII and NheI in plasmid pyEOE-attP-EXP2-3xMYC (14), replacing the *hsp86* promoter and *exp2* coding sequence. The mRuby3 coding sequence was then PCR amplified from pLN-HSP101-SP-mRuby3 (21) using primers P76/P77 and inserted between NheI and EagI, replacing the 3xMYC sequence and resulting in the plasmid pyEOE-attP-EXP2-5’UTR-SP-mRuby3. This plasmid was co-transfected with pINT (37) into EXP1^apt^ parasites, selection was applied 24 hours post-transfection with 2µM DSM1 and parasites were cloned upon returning from selection.

To monitor EXP2 by live fluorescence in EXP1^apt^, an endogenous mNeonGreen fusion to EXP2 was generated by co-transfecting EXP1^apt^ with plasmids pyPM2GT-EXP2-mNeonGreen (29) (linearized at AflII) and pUF-Cas9-EXP2-CT-gRNA (14). Selection was applied 24 hours post-transfection with 2µM DSM1 and parasites were cloned upon returning from selection.

### Proximity labeling and mass spectrometry

Parasites were synchronized to an ∼8 hour window by treatment with 5% w/v D-sorbitol and 200 µM exogenous biotin was added in the ring stage. After 18 hours, trophozoite and schizont-infected RBCs were purified on an LD column mounted on a QuadroMACs magnetic separator (Miltenyi Biotech), washed in PBS to remove residual biotin and lysed in RIPA buffer containing protease inhibitors. Lysates were briefly sonicated and insoluble material and hemozoin was cleared by centrifugation before loading onto streptavidin magnetic beads (Pierce). After rotating at 4°C overnight, beads were washed 5X with RIPA followed by 5X washes in 50 mM Tris 6.8 containing 8M urea.

Protein samples were reduced and alkylated using 5mM Tris (2-carboxyethyl) phosphine and 10mM iodoacetamide, respectively, and then enzymatically digested by the sequential addition of trypsin and lys-C proteases as described (38, 39). The digested peptides were desalted using Pierce C18 tips (Thermo Fisher Scientific), dried and resuspended in 5% formic acid. Approximately 1 µg of digested peptides were loaded onto a 25 cm long, 75 um inner diameter fused silica capillary packed in-house with bulk C18 reversed phase resin (1.9 um, 100A pores, Dr. Maisch GmbH). The 140-minute water-acetonitrile gradient was delivered using a Dionex Ultimate 3000 UHPLC system (Thermo Fisher Scientific) at a flow rate of 200 nl/min (Buffer A: water with 3% DMSO and 0.1% formic acid and Buffer B: acetonitrile with 3% DMSO and 0.1% formic acid). Eluted peptides were subsequently ionized by the application of a distal 2.2kv and introduced into the Orbitrap Fusion Lumos mass spectrometer (Thermo Fisher Scientific) and analyzed by tandem mass spectrometry (MS/MS). Data was acquired using a Data-Dependent Acquisition (DDA) method consisting of a full MS1 scan (Resolution = 120,000) followed by sequential MS2 scans (Resolution = 15,000) to utilize the remainder of the 3 second cycle time.

Data analysis was accomplished using the Integrated Proteomics pipeline 2 (Integrated Proteomics Applications, San Diego, CA). Data was searched against the protein database from *Plasmodium falciparum* 3D7 downloaded from UniprotKB (10,826 entries) on October 2013. MS/MS spectra searched using the ProLuCID algorithm followed by filtering of peptide-to-spectrum matches (PSMs) by DTASelect using a decoy database-estimated false discovery rate of <1%.

### Antibodies

The following antibodies were used for immunofluorescence assays (IFA) and western blot (WB) analysis at the indicated dilutions: mouse anti-HA monoclonal antibody HA.11 (Covance; 1:500 WB); rabbit polyclonal anti-HA SG77 (ThermoFisher; 1:500 IFA and WB); mouse anti-Flag monoclonal antibody clone M2 (Sigma; 1:300 IFA); mouse anti-EXP1 monoclonal antibody (20) (1:500 WB); mouse anti-EXP2 monoclonal antibody clone 7.7 (40) (1:500 IFA and WB); rabbit polyclonal anti-SBP1 (41) (1:500 IFA); mouse anti-cMYC monoclonal antibody 9E10 (ThermoFisher; 1:166 IFA and WB); rabbit polyclonal anti-*Plasmodium* Aldolase ab207494 (Abcam; 1:500 WB).

### Western blot

Western blots were carried out as previously described (14) and imaging with an Odyssey infrared imaging system (Li-COR Biosciences). Biotinylated proteins were detected with IRDye 800-conjugated streptavidin used at 1:200. Signal quantification was performed with Image Studio software (Li-COR Biosciences).

### Parasite growth assays

Parasite cultures were washed five times to remove aTc, then plated at 5% parasitemia (percentage of total RBCs infected) with or without 1µM aTc in triplicate. Every 24 hours, media was changed and parasitemia was measured by flow cytometry on an Attune NxT (ThermoFisher) by nucleic-acid staining with PBS containing 0.8 μg/ml acridine orange. Subculture (1:1) was performed each day parasitemia exceeded 10% and half as often for -aTc cultures within that group containing parasitemia less than 1%. Cumulative parasitemia was back-calculated based on subculture schedule, data were log2 transformed and fitted to a linear equation to determine slope using Prism (Graphpad).

### Transmission electron microscopy

EXP1^apt^ parasites were extensively washed to remove aTc, replated and allowed to develop 48 hours with or without aTc along with the parental NF54^attB^:HSP101-3xFLAG line. Trophozoite- and schizont-infected RBCs were purified on an LD column mounted on a QuadroMACs magnetic separator and fixed in 100 mM sodium cacodylate buffer, pH 7.2 containing 2% paraformaldehyde and 2.5% glutaraldehyde (Polysciences) for 1 hour at room temperature for ultrastructural analyses. Samples were washed in sodium cacodylate buffer at room temperature and post-fixed in 1% osmium tetroxide (Polysciences) for 1 hour. Samples were then extensively rinsed in dH20 before en bloc staining with 1% aqueous uranyl acetate (Ted Pella) for 1 h. Following several rinses in water, samples were dehydrated in a graded series of ethanol and embedded in Eponate 12 resin (Ted Pella). Sections (95 nm thick) were cut with a Leica Ultracut UCT ultramicrotome (Leica Microsystems), stained with uranyl acetate and lead citrate, and viewed on a JEOL 1200 EX transmission electron microscope (JEOL USA) equipped with an AMT 8-megapixel digital camera and AMT Image Capture Engine V602 software (Advanced Microscopy Techniques). For quantification of abnormalities, 100 trophozoites and 100 segmented schizonts were scored in each of two independent replicates.

### Immunofluorescence imaging

For IFAs, cells were fixed with a mixture of cold 90% acetone and 10% methanol for 2 minutes, except for export assays where fixation was carried out with room temperature 100% acetone for 2 minutes, and processed as described (9). For detection of biotinylated proteins, streptavidin-conjugated Alexa Fluor 594 (ThermoFisher) was included with secondary antibodies at 1:200. Images were collected on an Axio Observer 7 equipped with an Axiocam 702 mono camera and Zen 2.6 Pro software (Zeiss) using the same exposure times for all images across sample groups and experimental replicates.

### Quantification of protein export

IFA analysis of protein export was performed as described (14) except that image quantification was carried out using the Image Analysis module in Zen 2.6 Pro (Zeiss). The border of each single-infected RBC was traced using the DIC channel as a reference and the PVM was marked using the “Segment by Global Thresholding” tool for the HSP101-3xFLAG-488 channel (low and high thresholds set at 600 and 16,383 respectively and the Fill Holes option enabled). The signal corresponding to exported SBP1 was determined by removing any SBP1 signal within the PVM from the total SBP1 signal in each cell. Individual Maurer’s clefts were identified using the “Dynamic Thresholding” tool for the SBP1-594 channel (smoothing set to 7, threshold set to - 500, minimum area set to 10 and the Watersheds option enabled with count set to 1) and removing puncta within the PVM boundary from the total SBP1 puncta within each cell.

### Live-fluorescent imaging of EXP2-mNeonGreen distribution

Parasite cultures were washed 5 times to remove aTc, then plated with or without 1µM aTc and cultured for 48 hours. Trophozoite and schizont-infected RBCs were magnet purified and stained with 2.5 µM BODIPY TR Ceramide (Thermo Fisher) for 15 minutes at 37°C, washed once with media and immediately imaged. For each replicate, 40 single-infected RBCs were selected moving top to bottom and left to right in each field to avoid bias. For each infected RBC, the circumference of the PVM was traced with the profile tool in Zen 2.6 Pro using the BODIPY TR Ceramide signal at the parasite periphery as a guide with the green channel turned off to blind the experimenter to the mNeonGreen (mNG) signal. The mNG fluorescent intensity along each PVM trace was then collected, mNG signal and distance along the trace were normalized and data were analyzed with custom R scripts (ver. 3.5.2). In one approach, peaks were designated as mNG fluorescent intensity that exceeded the mean of the minimum and maximum intensity for at least 3 measured points before falling below this threshold. In an alternative approach, minimum PVM circumference containing indicated amounts of EXP2-mNG fluorescent signal was determined as the shortest distance along the PVM trace that returned the designated portion of total fluorescent signal throughout the trace. Means from three independent experiments were fitted to a smooth line with JMP Pro (ver. 14.2.0).

### Live-fluorescent imaging of PV-mRuby3

Parasite cultures were washed 5 times to remove aTc, then plated with or without 1µM aTc and cultured for 48 hours. Trophozoite and schizont-infected RBCs were magnet purified and stained with 1µg/ml Hoechst 33342 trihydrochloride trihydrate (ThermoFisher) for 15 min at 37°C, washed once with media and immediately imaged.

## Data Availability

Custom R scripts used in this study are available at https://github.com/tnessel/Beck-Lab

## Results

### Identification of proteins at the luminal face of the PVM with BioID2

To probe the protein content of the PV/PVM, we attempted to fuse the BioID proximity labeling system to the endogenous EXP2 protein. However, repeated attempts to generate an EXP2-BioID fusion with a verified CRISPR/Cas9 strategy for editing the 3’ end of *exp2* were unsuccessful. Endogenous EXP2 can tolerate a monomeric NeonGreen (mNG, 27 kDa) fusion without an obvious fitness cost (29); thus the bulkier size (35 kDa) or enzymatic activity of BioID may interfere with EXP2 trafficking or essential functions. To test the former possibility, we next attempted fusion with the second generation BioID2 derived from *Aquifex aeolicus* (Figure 1A). BioID2 lacks the DNA-binding domain present in the *E. coli*-derived BioID, resulting in a smaller size (27 kDa) that has been shown to reduce trafficking defects relative to BioID fusions (28).

**Figure 1:**
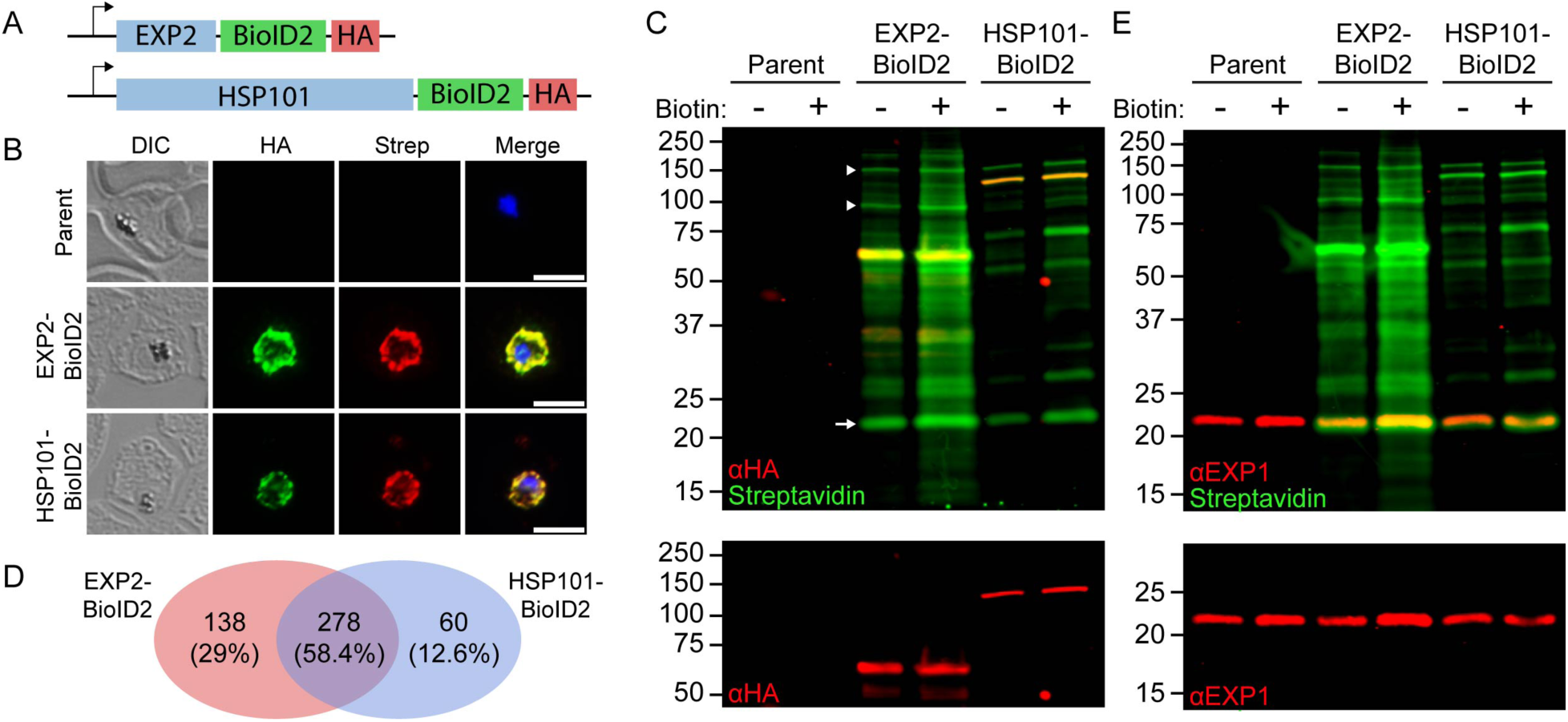
BioID2 identifies the protein contents of the PV/PVM. (A) Schematic showing C-terminal BioID2-3xHA fusion to the endogenous *exp2* and *hsp101* genes. (B) Immunofluorescence assay of parental, EXP2-BioID2 and HSP101-BioID2 parasites grown in media supplemented with 200 µM biotin. Scale bars are 5 µm. (C) Western blot of parental, EXP2-BioID2 and HSP101-BioID2 lines grown in regular RPMI (which contains 820 nM biotin, designated “-”) or 18 hours in RPMI supplemented with 200 µM biotin (designated “+”). Molecular weights after signal peptide cleavage are predicted to be 61.2 kDa for EXP2-BioID2-3xHA and 130 kDa for HSP101-BioID2-3xHA. Arrowheads from top to bottom indicate bands expected to correspond to untagged PTEX150 and HSP101. Arrow indicates prominent band at ∼23 kDa. (D) Ven diagram summarizing overlapping and distinct proteins detected in EXP2-BioID2 and HSP101-BioID2 datasets. All proteins identified in untagged negative controls were removed from the BioID2 datasets and remaining proteins that were present in both independent replicates of EXP2-BioID2 or HSP101-BioID2 were used to generate the diagram. (E) Western blot as in (C) probed with anti-EXP1 to show correspondence with prominent ∼23 kDa band. Molecular weight after signal peptide cleavage is predicted to be 14.7 kDa for EXP1. Note that EXP1 is observed to migrate at a higher molecular weight than predicted.

Parasites with an endogenous EXP2-BioID2 fusion were easily obtained and displayed robust biotinylation activity at the PVM (Figure S1A and Figure 1B), suggesting the additional ∼9 kDa BioID DNA-binding domain is not compatible with EXP2 function. As an additional probe, we also generated parasites with an endogenous HSP101-BioID2 fusion (Figure 1A,B). As EXP2 appears to be expressed at a stoichiometrically higher level than HSP101 and a fraction of PTEX-independent EXP2 exists (14, 24, 26), we reasoned that these two fusions would similarly position BioID2 within PTEX proximal to the luminal face of the PVM but might allow distinct proximity labeling by the PTEX-independent fraction of EXP2 that could provide clues to its dual function or organization in the PVM.

Western blot of EXP2-BioID2 and HSP101-BioID2 lysates showed extensive protein biotinylation relative to parental controls and supplementation of cultures with 200 µM exogenous biotin for 18 hours produced an approximately 2-fold increase in streptavidin signal over parallel cultures maintained in standard RPMI containing 820 nM biotin (Figure 1C).

Overall biotinylation levels were higher in EXP2-BioID2 lines which may reflect the higher level of EXP2 expression relative to HSP101 (14). Although streptavidin-labeled banding patterns were distinct between the EXP2-BioID2 and HSP101-BioID2 lysates, both lines showed several strongly labeled bands migrating at molecular weights consistent with other PTEX components (Figure 1C, arrowheads). As expected, the most strongly labeled band in each lysate corresponded with the BioID2-3xHA fusion. Notably, a band migrating at ∼23 kDa displayed prominent labeling in both lines (Figure 1C, arrow).

To identify labeled proteins, synchronized parasite cultures were supplemented with 200 µM biotin in the ring stage and allowed to develop for 18 hours. Parasite-infected RBCs were magnetically purified before lysis, streptavidin chromatography and subsequent analysis by mass spectrometry to identify biotinylated proteins. Mass spectrometry datasets from two independent experiments performed with EXP2-BioID2 or HSP101-BioID2 identified ∼10-fold more proteins than untagged parental controls and were substantially overlapping and highly enriched for known PV/PVM proteins (Figure 1D and Table S2). The highest-ranking proteins by normalized spectral abundance factor (NSAF) identified in both EXP2-BioID2 and HSP101-BioID2 included protein export machinery (PTEX and EPIC (42) complexes) and other proteins known to reside in the PV lumen, exported proteins and the PVM membrane proteins EXP1 and ETRAMP10.2 (Table 1). Beyond the top ∼10% of each dataset, many additional known PV/PVM and exported proteins were identified (Table S2).

**TABLE 1.** Top 30 proteins identified in EXP2-BioID2 and HSP101-BioID2 experiments ranked by NSAF score.

| # | EXP2-BioID2_replicate 1 |  | EXP2-BioID2_replicate 2 |  | HSP101-BioID2_replicate 1 |  | HSP101-BioID2_replicate 2 |  |
| --- | --- | --- | --- | --- | --- | --- | --- | --- |
|  | Gene ID | Name or localization | Gene ID | Name or localization | Gene ID | Name or localization | Gene ID | Name or localization |
| 1 | PF3D7_1471100 | EXP2 | PF3D7_1436300 | PTEX150 | PF3D7_1436300 | PTEX150 | PF3D7_1116800 | HSP101 |
| 2 | PF3D7_1121600 | EXP1 | PF3D7_1033200 | ETRAMP10.2 | PF3D7_1116800 | HSP101 | PF3D7_1436300 | PTEX150 |
| 3 | PF3D7_1436300 | PTEX150 | PF3D7_1135400 | PV Localization (ref 40) | PF3D7_0721100 | None | PF3D7_1212000 | TPx(GI) |
| 4 | PF3D7_1135400 | PV Localization (ref 40) | PF3D7_1471100 | EXP2 | PF3D7_1212000 | TPx(GI) | PF3D7_0721100 | None |
| 5 | PF3D7_1033200 | ETRAMP10.2 | PF3D7_1024800 | EXP3 | PF3D7_1033200 | ETRAMP10.2 | PF3D7_1226900 | PV2 |
| 6 | PF3D7_1335100 | MSP7 | PF3D7_1212000 | TPx(GI) | PF3D7_1135400 | PVLocalization (ref 40) | PF3D7_1121600 | EXP1 |
| 7 | PF3D7_1226900 | PV2 | PF3D7_1129100 | PV1 | PF3D7_0702500 | Exported Protein | PF3D7_1108600 | ERC |
| 8 | PF3D7_0207500 | SERA6 | PF3D7_0207500 | SERA6 | PF3D7_1024800 | EXP3 | PF3D7_1471100 | EXP2 |
| 9 | PF3D7_1129100 | PV1 | PF3D7_1335100 | MSP7 | PF3D7_1121600 | EXP1 | PF3D7_1135400 | PV Localization (ref 40) |
| 10 | PF3D7_1212000 | TPx(GI) | PF3D7_1226900 | PV2 | PF3D7_1226900 | PV2 | PF3D7_0801000 | PHISTc |
| 11 | PF3D7_0721100 | None | PF3D7_1345100 | TRX2 | PF3D7_1108600 | ERC | PF3D7_1033200 | ETRAMP10.2 |
| 12 | PF3D7_1312700 | None | PF3D7_1121600 | EXP1 | PF3D7_1129100 | PV1 | PF3D7_1129100 | PV1 |
| 13 | PF3D7_1116800 | HSP101 | PF3D7_0532100 | ETRAMP5 | PF3D7_0801000 | PHISTc | PF3D7_1130100 | RPL38 |
| 14 | PF3D7_1024800 | EXP3 | PF3D7_1105600 | PTEX88 | PF3D7_0629200 | Partial PV Localization (ref 40) | PF3D7_0702500 | Exported Protein |
| 15 | PF3D7_1105600 | PTEX88 | PF3D7_0612700 | PTEX88 | PF3D7_1130100 | RPL38 | PF3D7_0625400 | None |
| 16 | PF3D7_0316300.1 | PPase | PF3D7_1116700 | DPAP1 | PF3D7_1471100 | EXP2 | PF3D7_1105600 | PTEX88 |
| 17 | PF3D7_1464600 | UIS2 | PF3D7_0316300.1 | PPase | PF3D7_1105600 | PTEX88 | PF3D7_0202400 | PTEF |
| 18 | PF3D7_1228600 | MSP9 | PF3D7_0721100 | None | PF3D7_1460700 | RPL27 | PF3D7_1404900 | None |
| 19 | PF3D7_0316300.2 | PPase | PF3D7_1116800 | HSP101 | PF3D7_0202400 | PTEF | PF3D7_0629200 | Partial PV Localization (ref 40) |
| 20 | PF3D7_0902800 | SERA9 | PF3D7_0404900 | P41 | PF3D7_1104400 | Trx-mero | PF3D7_1345100 | TRX2 |
| 21 | PF3D7_1345100 | TRX2 | PF3D7_1464600 | UIS2 | PF3D7_1345100 | TRX2 | PF3D7_1108700 | Pfj2 |
| 22 | PF3D7_1419800 | GR | PF3D7_0316300.2 | PPase | PF3D7_1404900 | None | PF3D7_1335100 | MSP7 |
| 23 | PF3D7_0207800 | SERA3 | PF3D7_1419800 | GR | PF3D7_1108700 | Pfj2 | PF3D7_1024800 | EXP3 |
| 24 | PF3D7_0532100 | ETRAMP5 | PF3D7_1334800 | MSRP2 | PF3D7_0830400 | PV/Exported Localization (ref 76) | PF3D7_1008900 | AK1 |
| 25 | PF3D7_1420700 | P113 | PF3D7_1454400 | APP | PF3D7_1016400 | FIKK10.1 | PF3D7_1232100 | CPN60 |
| 26 | PF3D7_1014100 | MSA180 | PF3D7_0830400 | PV/Exported Localization (ref 76) | PF3D7_1232100 | CPN60 | PF3D7_0706400 | RPL37 |
| 27 | PF3D7_0207400 | SERA7 | PF3D7_0902800 | SERA9 | PF3D7_1008900 | AK1 | PF3D7_1104400 | Trx-mero |
| 28 | PF3D7_0702500 | Exported | PF3D7_0207800 | SERA3 | PF3D7_1335100 | MSP7 | PF3D7_1016400 | FIKK10.1 |
| 29 | PF3D7_0501200 | PIESP2 | PF3D7_0930300 | MSP1 | PF3D7_0625400 | None | PF3D7_0501200 | PIESP2 |
| 30 | PF3D7_1419800 | GR | PF3D7_0207400 | SERA7 | PF3D7_1001200 | ACBP2 | PF3D7_1460700 | RPL27 |

**Table 2.**
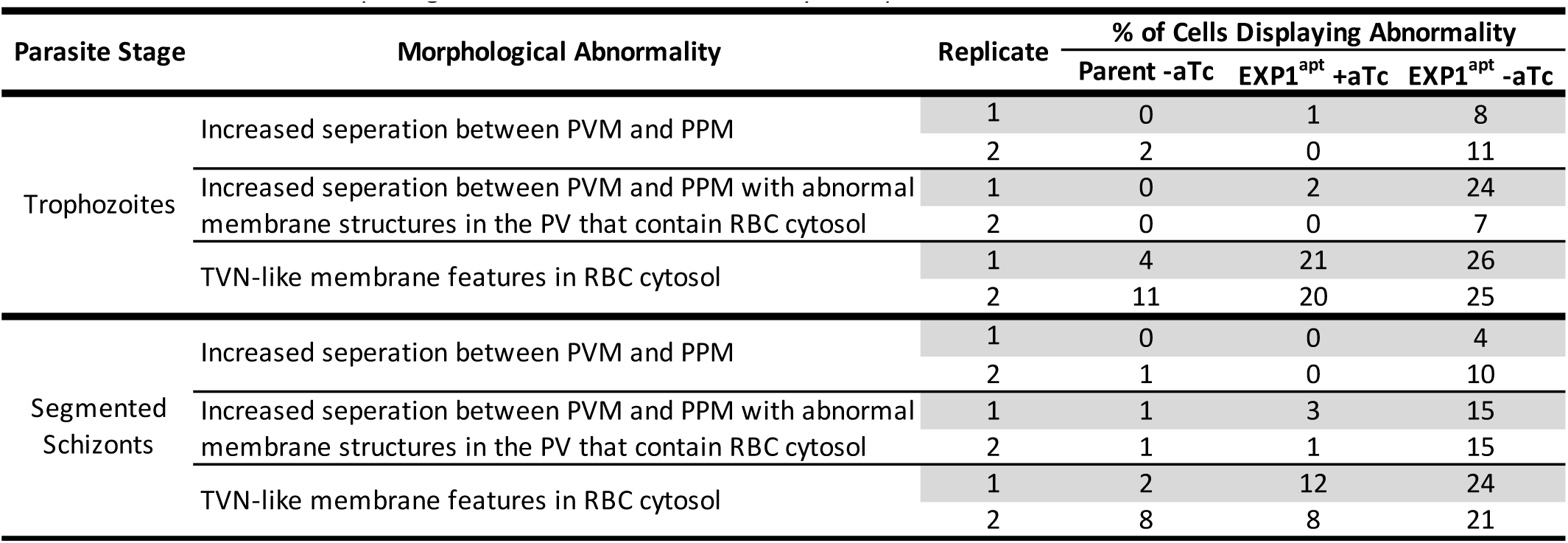
Quantification of morphological abnormalities in TEM analysis of parent and EXP1^apt^ lines.

EXP1 and ETRAMP10.2, both single-pass PVM membrane proteins with similar expression timing to EXP2 (Figure S1B), ranked particularly high across all BioID2 experiments. When Western blots of lysates from EXP2-BioID2 and HSP101-BioID2 were probed with an anti-EXP1 monoclonal antibody, the signal coincided with the prominent ∼23 kDa band labeled by streptavidin (EXP1 is known to migrate at 23 kDa, higher than its predicated molecular weight of 14.7 kDa following signal peptide cleavage (43)), consistent with the high level of EXP1 representation in the BioID2 proteomics (Figure 1E). ETRAMP5, another member of this group, was also well represented (though not consistently among the top hits). The functions of ETRAMP10.2, ETRAMP5 and EXP1 are not known. Given their high level of representation implying particular abundance and/or an intimate proximity to EXP2/PTEX, we focused on functional characterization of these three proteins.

### Efficient genome editing in *P. falciparum* with Cpf1/Cas12a

CRISPR/Cas9 genome editing technology has been rapidly adapted for manipulation of *P. falciparum*. A limitation of Cas9-mediated editing is the requirement for a “GG” in the protospacer adjustment motif (PAM), as such sites are comparatively rare given the high A+T content of the *P. falciparum* genome (80.6%) (44). The type II, class V CRISPR endonuclease Cpf1 (CRISPR from *Prevotella* and *Francisella*, also known as Cas12a) has recently emerged as a favorable alternative to Cas9 with an inherently lower level of off-target cleavage (45, 46). In contrast to Cas9, Cpf1 does not require a trans activating CRISPR (tracr) RNA, introduces a staggered double-strand break in target DNA that is distal from the PAM and utilizes a T-rich PAM, making it uniquely suited to editing the *P. falciparum* genome (32).

Cpf1 from *Acidaminococcus sp. BV3L6* and *Lachnospiraceae bacterium ND2006* (AsCpf1 and LbCpf1, respectively) have been shown to mediate efficient genome editing in mammalian cells (32) and we tested both of these enzymes for their genome editing capacity in *P. falciparum* relative to a Cas9 editing system we previously developed (14, 31). The 3’ region of *exp1* was found to contain attractive Cas9 and Cpf1 target sites located in close proximity, providing an opportunity for initial Cpf1 testing (Figure 2A). We generated plasmids for expression of AsCpf1 or LbCpf1 and corresponding gRNAs to target two sites with Cpf1 near the *exp1* stop codon. In parallel, we generated constructs for targeting partially overlapping or immediately adjacent sites with Cas9 (Figure 2A). These markerless Cas9 and Cpf1 editing plasmids were co-transfected with a donor plasmid bearing a yDHODH cassette and flanks designed to repair the intended double-strand breaks by double homologous recombination to introduce a 3xHA-GFP11 tag at the 3’ end of *exp1* (Figure 2A). In each case, parasites returned from DSM1 selection in about 21 days with the expected edit, as gauged by diagnostic PCR and Western blot (Figure 2B). To provide a tool for marker-free editing, we additionally generated AsCpf1 and LbCpf1 vectors with a yDHODH cassette to allow for selection in parasites and showed that these vectors also facilitated editing of the *exp1* locus (Figure S2A).

**Figure 2:**
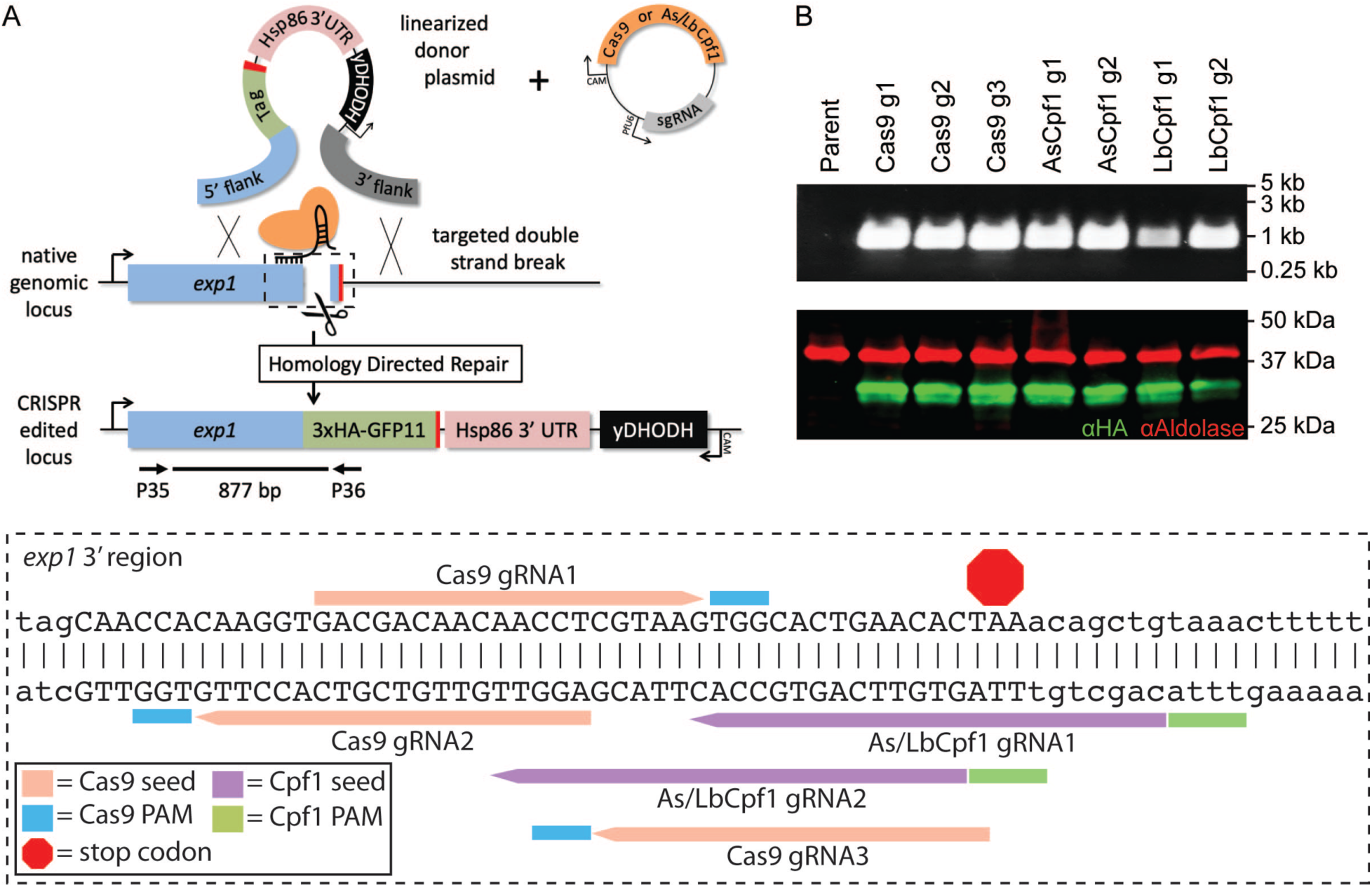
Genome editing of the *exp1* locus with Cpf1. (A) Schematic showing strategy for double homologous recombination repair of double-strand breaks mediated by Cas9 or Cpf1 at the 3’ end of *exp1.* Individual gRNA target sequences are shown in the zoomed view in the dashed box. Exon sequences are shown in uppercase while intron and UTR sequences are shown in lowercase. 3’ UTR, 3’ untranslated region; yDHODH, yeast dihydroorotate dehydrogenase; PAM, protospacer adjustment motif. (B) Diagnostic PCR with primers indicated in the schematic and Western blot showing successful integration at the 3’ end of *exp1* mediated by either Cas9 or Cpf1. Aldolase serves as a loading control. Molecular weight after signal peptide cleavage is predicted to be 20.9 kDa for EXP1-3xHA-GFP11. Note that EXP1 and derivative fusions are observed to migrate at a higher molecular weight than predicted.

Encouraged by these results, we next applied the AsCpf1 system to edit the *etramp10.2* and *etramp5* loci to insert a 3xHA-epitope and simultaneously install the TetR-DOZI-aptamers (TDA) system for conditional translational repression (Figure S2B-D) (31, 33). Editing of the 3’ end of both loci was also successful (Figure S2C,D), however both ETRAMP10.2^apt^ and ETRAMP5^apt^ parasite lines were found to a have truncated 3’ aptamer arrays (reduced from 10X to 6X, data not shown). Partial truncation of 3’ aptamers rapidly diminishes translational control (31) and anhydrotetracycline (aTc) washout did not result in measurable knockdown (Figure S2C,D), thus ETRAMP10.2 and ETRAMP5 were not pursued further in this study. Collectively, these results indicate Cpf1 editing of the *P. falciparum* genome is equally efficient to Cas9 with Cpf1 offering many additional gRNA targets owing to its T-rich PAM requirement.

### Lethal EXP1 knockdown with a dual aptamer strategy

To query EXP1 function in PV biology, we employed a dual aptamer TDA strategy using a linear plasmid system to replace the endogenous *exp1* coding sequence in an NF54^attB^ parasite line bearing a 3xFLAG tag on the endogenous *hsp101* gene (10, 14). This was accomplished by LbCpf1 editing to introduce an aptamer just upstream of the start codon and a 10X aptamer array just downstream of the stop codon (Figure 3A and Figure S3A,B). Installation of 5’ aptamers has been shown to reduced baseline expression even in the presence of aTc (14) and the resulting EXP1^apt^ parasites showed 87.5±7.2% reduction in EXP1 expression and a growth defect relative to the parent line (Figure 3B,C). Removal of aTc further reduced EXP1 levels (99.3±0.4% knockdown relative to parent by probing with anti-EXP1 and 65.9±3.1% relative to EXP1^apt^ +aTc by probing with anti-HA) and resulted in a complete block in parasite growth, indicating EXP1 is required for intraerythrocytic development (Figure 3B,C).

**Figure 3:**
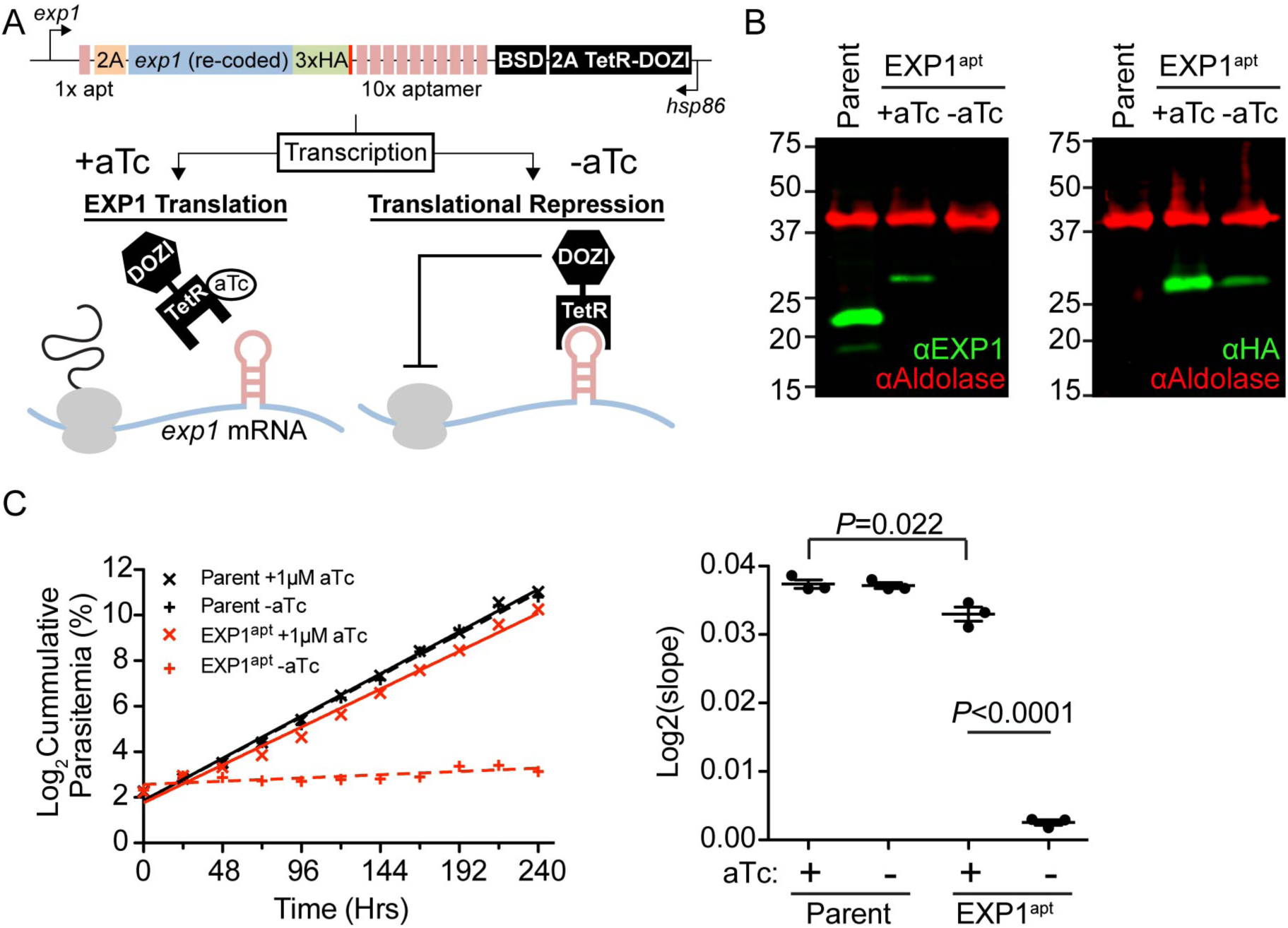
Lethal EXP1 knockdown with a dual aptamer strategy. (A) Schematic of modified *exp1* locus and TDA control of translation in EXP1^apt^ parasites. 2A, *Thosea asigna* virus 2A skip peptide; BSD, blasticidin-S deaminase. (B) Western blot of parental parasite line grown without aTc and EXP1^apt^ parasites grown 48 hours with or without aTc. Aldolase serves as a loading control. Molecular weights after signal peptide cleavage are predicted to be 14.7 kDa for EXP1 and 18 kDa for EXP1-3xHA. Note that EXP1 and derivative fusions are observed to migrate at a higher molecular weight than predicted. Results are representative of three independent experiments. (C) Growth analysis of parental and EXP1^apt^ parasites with or without aTc. Results from one experiment with three technical replicates plotted are shown. The scatter plot shows the slope of the line fitted to the mean of log2-transformed parasitemias for each of three independent experiments. Error bars indicate SEM. P values were determined by an unpaired, two-sided Student’s t-test.

As a complement to the titratable translational control afforded by the TDA system and to simultaneously provide a parallel option for conditional knockout, we also engineered the donor plasmid to place *loxP* sites around the recoded *exp1* gene and inserted cassettes for expression of the rapamycin-inducible dimerizable Cre recombinase (DiCre) downstream of the modified *exp1* locus (Figure S3A). Treatment of EXP1^apt^ cultures with rapamycin induced the expected excision event as gauged by diagnostic PCR but produced only modest impact on parasite growth (Figure S2C-E). Consistent with this, the non-excised locus remained readily detectable in these cultures even when parasites were grown for several days with rapamycin, indicating inefficient excision of *exp1* that was unsuitable for functional analysis (Figure S2E). The reason for the poor excision efficiency by DiCre in EXP1^apt^ was not explored further and the robust translational knockdown achieved by TDA was used for the remainder of the study.

### EXP1 functional constraints

While a 3xHA-GFP11 tag (6.2 kDa) could be efficiently fused to the endogenous EXP1 C-terminus (Figure 2), multiple attempts using the same editing strategy failed to generate an endogenous fusion to mNG or BioID2, suggesting the essential function of EXP1 is perturbed by bulky C-terminal fusions (data not shown). To directly test this possibility, we complemented the EXP1^apt^ parasites with a second copy of EXP1 bearing either a 3xMYC (4.5 kDa) or mNG-3xMYC (31.5 kDa) fusion (Figure 4A). While both versions of EXP1 were similar expressed, only the 3xMYC fusion was able to rescue parasite growth upon aTc removal (percent growth rate in -aTc relative to +aTc control was 7.28±20% in mNG-3xMYC compared with 67.63±7.36% in WT-3xMYC and 7.76±2.14% in uncomplemented EXP1^apt^), indicating that introduction of a bulky C-terminal fusion does indeed ablate EXP1 function (Figure 4B,C). Similar to EXP1-3xMYC, EXP1-mNG-3xMYC still localized at the parasite periphery with endogenous EXP1-3xHA but tended to show a more dispersed distribution possibly indicating perturbations in trafficking (Figure 4D).

**Figure 4:**
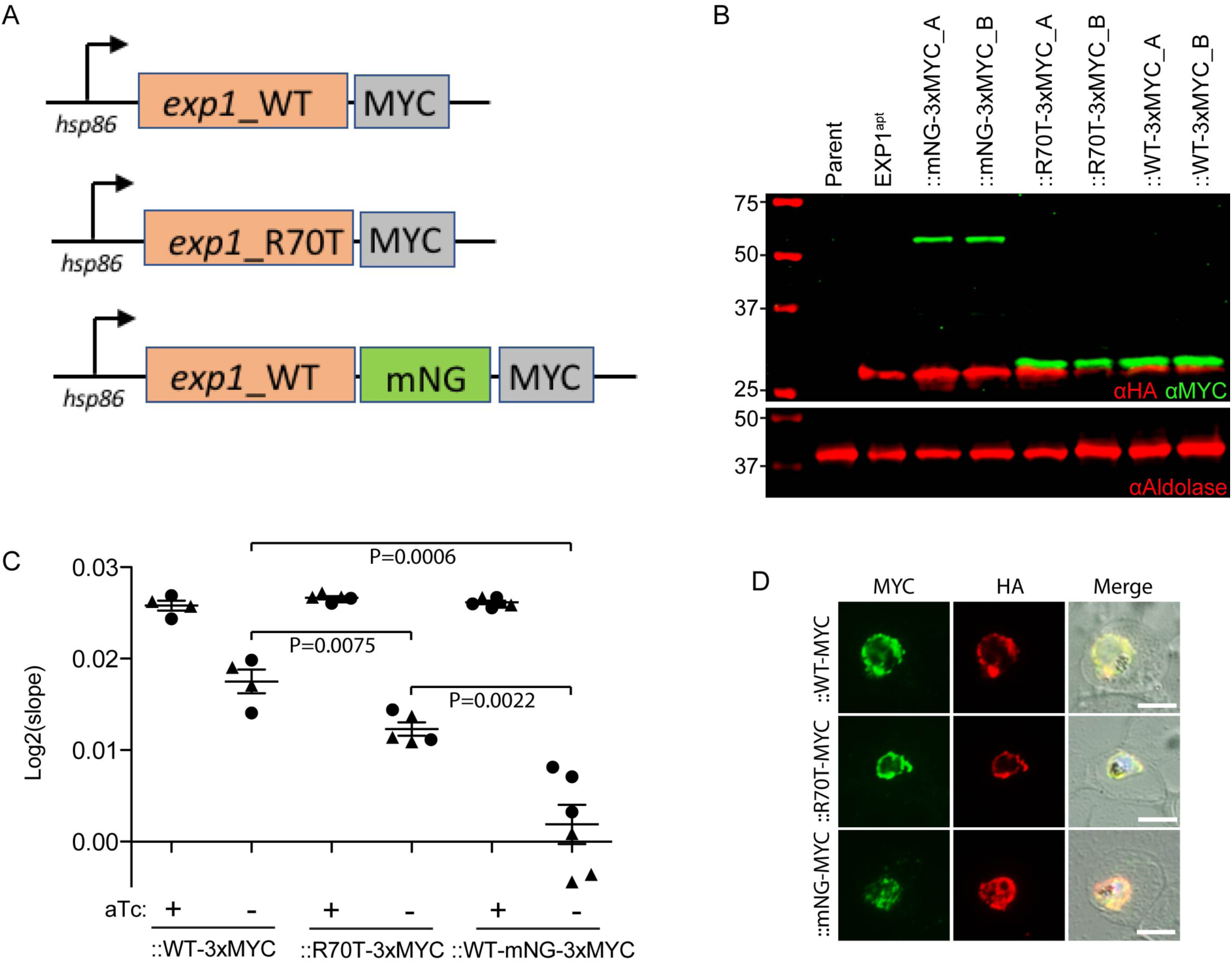
EXP1 *in vivo* function is ablated by a bulky C-terminal fusion but not by mutation of a residue important for GST activity *in vitro*. (A) Schematic showing complementing versions of EXP1 introduced into EXP1^apt^. (B) Western blot of parent, EXP1^apt^ and complemented lines. Two independently complemented lines were generated with each construct and are designated A or B. Aldolase serves as a loading control. Molecular weights after signal peptide cleavage are predicted to be 18 kDa for EXP1-3xHA, 19.2 kDa for EXP1-WT-3xMYC and EXP1-R70T-3xMYC and 46.2 kDa for EXP1-mNG-3xMYC. Note that EXP1 and derivative fusions are observed to migrate at a higher molecular weight than predicted. (C) Growth analysis of complemented EXP1^apt^ lines with or without aTc. The scatter plot shows the slope of the line fitted to the mean of log2-transformed parasitemias for four (EXP1-WT-3xMYC), five (EXP1-R70T-3xMYC) or six (EXP1-mNG-3xMYC) independent experiments. Means from independently generated lines complemented with the same version of EXP1 were pooled and are distinguished by different symbols (circles and triangles). Error bars indicate SEM. P values were determined by an unpaired, two-sided Student’s t-test. (D) Immunofluorescence assay of EXP1^apt^ complemented lines. Scale bars are 5 µm.

EXP1 has previously been reported to possess GST activity based on an *in silco* functional prediction approach and *in vitro* biochemical analysis (20). This study identified the arginine at position 70 in EXP1 as critical for GST activity *in vitro* and our functional complementation system provided the opportunity to test the importance of this residue *in vivo* by complementing EXP1^apt^ parasites with an EXP1-R70T-3xMYC mutant (Figure 4A,B). Similar to the wild-type second copy, EXP1-R70T-3xMYC colocalized with endogenous EXP1-3xHA at the PVM (Figure 4D). To our surprise, the R70T mutant also provided substantial rescue upon knockdown of endogenous EXP1 (percent growth rate in -aTc relative to +aTc control was 46.21±6.05% in R70T-3xMYC compared with 67.63±7.36% in WT-3xMYC), suggesting that GST activity cannot fully explain the *in vivo* function of EXP1 (Figure 4C).

### Depletion of EXP1 results in late cycle arrest and PV/PVM morphological abnormalities

While EXP1 expression peaks about midway through the intraerythrocytic development cycle (Figure S1B), EXP1 has been localized to merozoite dense granules and is thus expected to be delivered to the PVM immediately following RBC invasion and accumulate to peak levels later in the cycle (47). To determine the impact of EXP1 knockdown on parasite development from the point of invasion on, trophozoites (∼32-42 hours post invasion) were magnet-purified from synchronized cultures, washed of aTc to prevent new synthesis of EXP1 during dense granule formation at the terminal stages of schizogony and used to initiate cultures with fresh uninfected RBCs. New ring formation was not impacted relative to controls maintained with aTc, indicating EXP1 is not important at this early stage; rather cultures developed normally until a late stage when parasites arrested predominantly as trophozoites and schizonts and failed to complete the cycle (Figure 5A). We compared these developmental defects with those following inactivation of protein export and PVM channel activity in EXP2^apt^, an EXP2 conditional mutant that we previously generated using the same dual aptamer TDA approach (14). EXP2^apt^ parasites grown without aTc arrested at an earlier trophozoite stage that was distinct from EXP1-depleted parasites, suggesting EXP2-dependent transport activities are not impacted by loss of EXP1 (Figure 5B).

**Figure 5:**
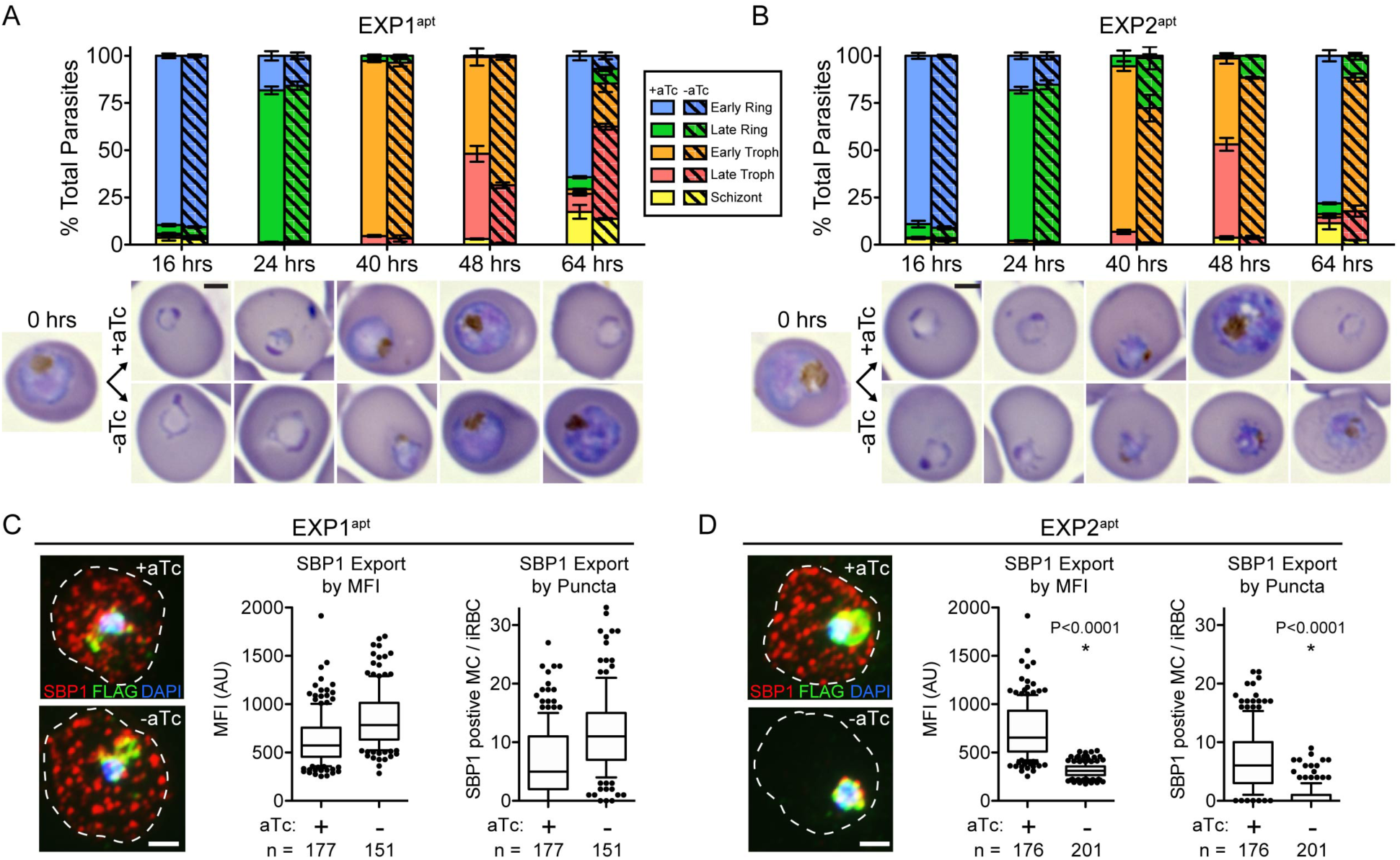
Depletion of EXP1 results in late cycle arrest but does not impact protein export. (A-B) Quantification of parasite stages of development from Giemsa-stained thin smears of (A) EXP1^apt^ and (B) EXP2^apt^ parasites synchronized to a 10-hour window and grown with or without aTc. Time 0 hours indicates the point of aTc removal at which purified late trophozoites (∼32-42 hours post-invasion) were mixed with fresh uninfected RBCs. Results from one experiment performed in technical triplicate are shown and are representative of two independent experiments. Error bars indicate SD. Representative images of the majority parasite population at each time point are shown. Scale bars are 2 µm. (C-D) Immunofluorescence assay showing SBP1, which is exported to the Maurer’s clefts, in (C) EXP1^apt^ and (D) EXP2^apt^ parasites. Quantification of SBP1 export beyond the PVM (both EXP1^apt^ and EXP2^apt^ contain a 3xFLAG tag on HSP101 which was used as a marker for the PVM) is shown as MFI of SBP1 within the host compartment or as the number of SBP1-positive puncta (Maurer’s clefts) within the host compartment. Merged images of SBP1, FLAG and DAPI signal are shown. The dashed line indicates the boundary of the host RBC traced from the corresponding DIC image. Data are pooled from three independent experiments and *n* is the number of individual parasite-infected RBCs. Boxes and whiskers delineate 25^th^-75^th^ and 10^th^-90^th^ percentiles, respectively. P values were determined by an unpaired, two-sided Student’s t-test. MFI, mean fluorescence intensity; AU, arbitrary units. Scale bars are 2 µm.

To directly test this, we monitored protein export beyond the PVM in EXP1^apt^ and EXP2^apt^ parasites using the above experimental design of removing aTc from synchronized, purified trophozoites and then analyzing export by IFA at the midpoint of the following cycle. While EXP2 knockdown results in a robust block in export of SBP1 as previously reported (14), no defect in SBP1 export was observed following knockdown of EXP1 (Figure 5C,D). Rather, SBP1 export was more robust in the absence of aTc as measured by mean fluorescent intensity in the infected RBC compartment or the number of SBP1-positive Mauer’s clefts (punctate structures beyond the PVM) per infected RBC (Figure 5C). While the basis for the apparent increase in exported SBP1 signal is unclear, these results clearly indicate EXP1 is not required for protein export.

To better understand the impact of EXP1 depletion in late stage parasites, we next examined parasite ultrastructure by transmission electron microscopy (TEM). As seen in the parental line and EXP1^apt^ grown with aTc, the PVM normally tightly overlays the PPM (Figure 6A,B). In contrast, EXP1^apt^ parasites grown 48 hours without aTc displayed striking changes in PV morphology with increased separation between the PVM and PPM in both trophozoites and segmented schizonts (Figure 6C-E). Curiously, the lumen of these enlarged PVs often contained additional membrane-enclosed structures filled with host cytosol (Figure 6D, double arrowheads). Additionally, membrane structures in the host cell cytosol external to the PVM (Figure 6B, enlarged region) which likely represent portions of the tubulovesicular network (TVN) were observed with increasing frequency upon EXP1 depletion. Quantification of morphological abnormalities observed by TEM in two independent experiments is shown in Table 2.

**Figure 6:**
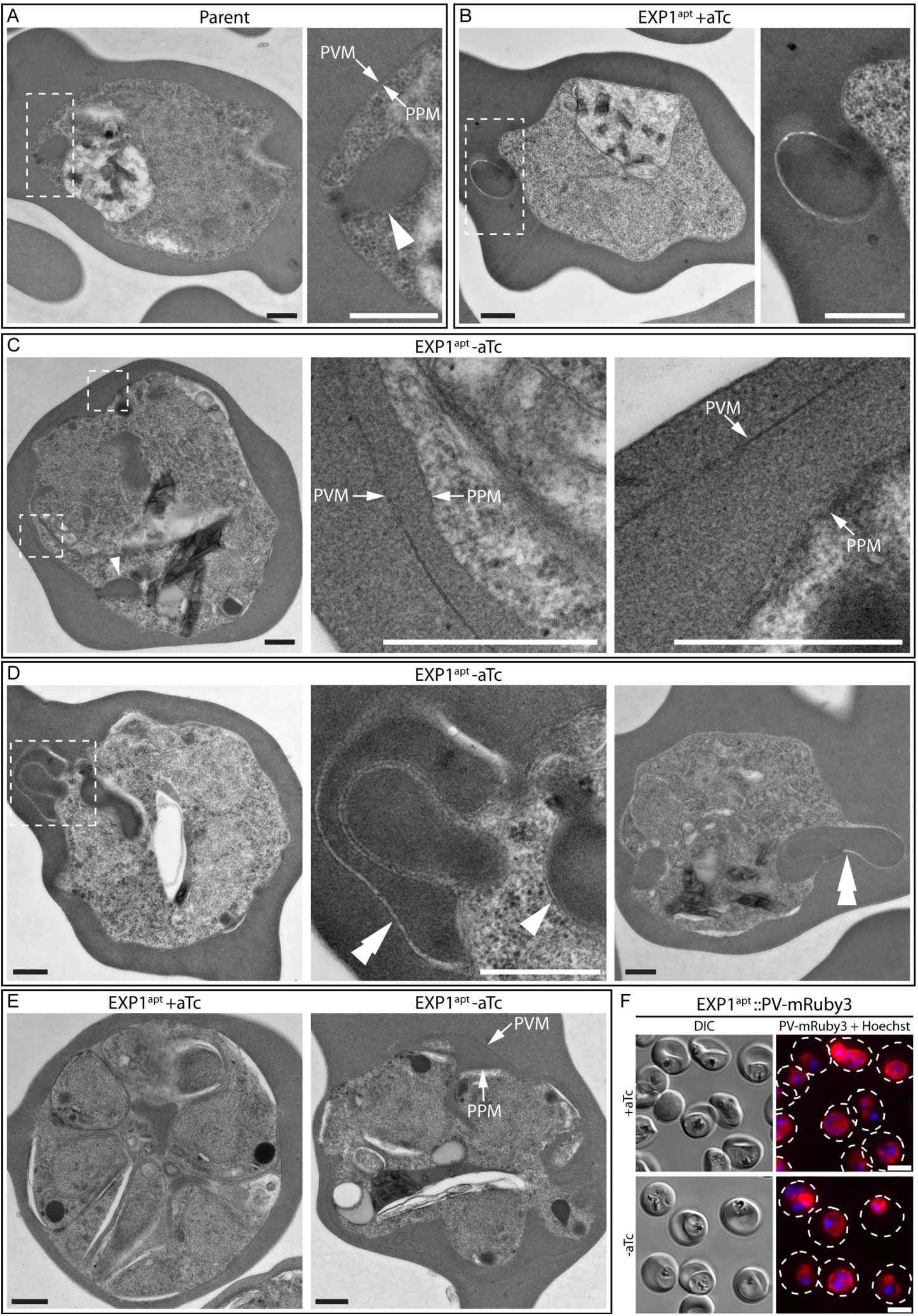
Depletion of EXP1 results in PV/PVM morphological abnormalities. (A-E) TEM visualization of parasite ultrastructure in parent and EXP1^apt^ parasites grown 48 hours with or without aTc. Images are shown of (A-D) trophozoites and (E) segmented schizonts. Dashed boxes indicate enlarged areas shown to the right. Arrows indicate PVM and PPM. Arrowheads indicate cytostomes. Double arrowheads indicate abnormal membrane-enclosed structures filled with host cytosol in the PV lumen. Results are representative of two independent experiments. Quantification of morphological abnormalities is shown in Table 2. Scale bars are 500 nm. (F) Live fluorescence imaging of magnet-purified EXP1^apt^::PV-mRuby3 parasites grown 48 hours with or without aTc. The dashed lines indicate the boundary of infected RBC traced from the corresponding DIC image. Results are representative of three independent experiments. Scale bars are 5 µm.

In EXP1^apt^ parasites that displayed increased separation between PVM and PPM, the enlarged PV lumen often showed equivalent density with the host cytosol, possibly indicating a broken vacuole (Figure 6C,E). To ascertain if PVM integrity was compromised following EXP1 knockdown, we fused a signal peptide to the fluorescent protein mRuby3 to target it to the PV and expressed it under the control of the *exp2* promoter in the EXP1^apt^ line. Parasites depleted of EXP1 continued to show concentrated mRuby3 signal at the cell periphery, indicating the PVM is not compromised, despite altered PVM ultrastructure (Figure 6F).

### EXP1 is required for proper organization of EXP2 in the PVM

To further investigate alterations in the PV, we evaluated the impact of EXP1 knockdown on other PVM proteins. As expected from the observation that protein export remains operational in EXP1^apt^, depletion of EXP1 did not substantially alter EXP2 levels (Figure 7A). To monitor EXP2 distribution in EXP1^apt^ parasites, we introduced a C-terminal mNG fusion on the endogenous copy of EXP2 and imaged parasites at a late stage corresponding with developmental arrest. Live fluorescent analysis of a parental EXP2-mNG control line with an unmodified *exp1* locus (29) showed a punctate distribution at early stages that often resolved into several larger patches in trophozoites and schizonts (Figure 7B). In contrast, EXP2 distribution was substantially altered in EXP1^apt^::EXP2-mNG parasites, often concentrating into one or two discrete points along the PVM (Figure 7B). To quantify this altered localization of EXP2-mNG, BODIPY-TR-Ceramide labeling was used as a guide to trace the PVM and collect EXP2-mNG signal along the PVM circumference (Figure 7B). Analysis of these PVM traces showed that the number of discrete EXP2 signal patches around the PVM was significantly reduced in EXP1^apt^ parasites with EXP2 signal concentrated into a smaller proportion of the PVM circumference (Figure 7C,D). These results show that although EXP2 function is preserved following depletion of EXP1, its organization in the PVM is drastically altered.

**Figure 7:**
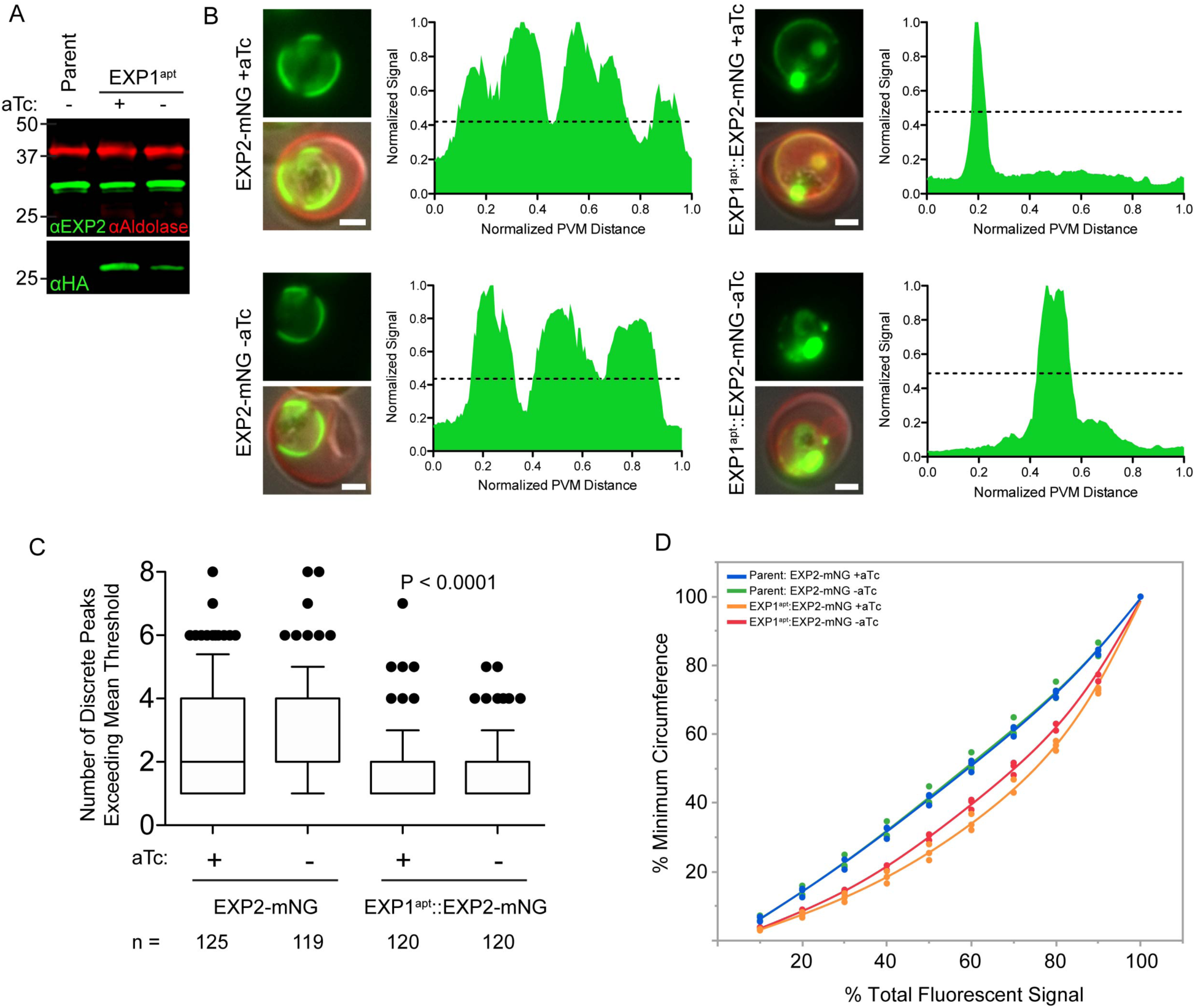
Depletion of EXP1 alters EXP2 distribution in the PVM. (A) Western blot of parental parasite line grown without aTc and EXP1^apt^ parasites grown 48 hours with or without aTc. Aldolase serves as a loading control. Molecular weights after signal peptide cleavage are predicted to be 30.8 kDa for EXP2 and 18 kDa for EXP1-3xHA. Note that EXP1 and derivative fusions are observed to migrate at a higher molecular weight than predicted. (B) Live fluorescent images of EXP2-mNG and EXP1^apt^::EXP2-mNG parasites grown 48 hours with or without aTc. Merged images include EXP2-mNG signal in green together with DIC and BODIPY TR Ceramide signal in red used to trace the PVM. Scale bars are 2 µm. The corresponding histograms of EXP2-mNG signal along the PVM trace are shown for each cell. Dashed line is the mean of the minimum and maximum EXP2-mNG signal intensity. (C) Quantification of the number of discrete peaks of EXP2-mNG signal exceeding the mean signal threshold as shown in (B) for EXP2-mNG and EXP1^apt^::EXP2-mNG parasites grown 48 hours with or without aTc. Data are pooled from three independent experiments and *n* is the number of individual parasites. Boxes and whiskers delineate 25^th^-75^th^ and 10^th^-90^th^ percentiles, respectively. P values were determined by an unpaired, two-sided Student’s t-test. (D) Graph showing the minimum distance along the PVM (given as a percent of total PVM length) containing the indicated amounts of EXP2-mNG fluorescent signal (given as percent of total EXP2-mNG signal per trace) in EXP2-mNG and EXP1^apt^::EXP2-mNG parasites grown 48 hours with or without aTc. Data points are means from three independent experiments fitted to a smooth line.

## Discussion

Promiscuous biotin ligases have emerged as powerful tools for proximity-based protein identification in live cells (48). To date, the original *E. coli* BirA* (BioID) has been employed by four studies in *Plasmodium spp*., including two that targeted BioID to the PV (49–52). Khosh-Naucke and colleagues targeted a signal peptide-GFP-BirA* fusion to the *P. falciparum* PV lumen (50) while Schnider and colleagues utilized a second copy of EXP1 with a C-terminal BirA* fusion to probe the PV in *P. berghei* (51). In the present study, we present the first use of BioID2 in *Plasmodium spp*., which we fused to the endogenous copy of EXP2 or HSP101 to identify proteins at the luminal face of the PVM. The ability to fuse mNG or BioID2 but not BioID to EXP2 indicates EXP2 will not tolerate larger bulky fusions, possibly due to defects in trafficking, oligomerization or assembly with other PTEX components, demonstrating that the smaller size of BioID2 offers advantages over the original BioID system. We have also adapted a Cpf1 editing system for use in *P. falciparum* which facilitated successful genome editing with the same efficiency as our Cas9 editing system. Since the discovery of its T-rich PAM requirements, Cpf1 has been suggested to possess unique promise for manipulation of the A+T rich genomes of *Plasmodium spp.* (32). To our knowledge, this is the first report of Cpf1 use in malaria parasites, realizing that potential of greatly expanding the repertoire of gRNA targets available to manipulate the *P. falciparum* genome.

While we expect the EXP2-BioID2 and HSP101-BioID2 datasets to contain novel PV and PVM proteins (Figure S1C) and are currently working to validate these candidates, we have focused in the present study on functional analysis of EXP1, one of the highest-ranking proteins in our proteomics. The study of EXP1 dates back more than 35 years when a monoclonal antibody shown to label the PVM (40, 53, 54) was subsequently found to detect an antigen in the EXP1 C-terminus (43). However, despite being one of the earliest discovered PVM proteins, EXP1 function has remained obscure. Previous failed attempts to disrupt the *exp1* gene in *P. falciparum* (55) and *P. berghei* (56) indicated a critical role for EXP1 during the blood stage.

EXP1 is also expressed and localized to the PVM in the parasite liver stage (57) and a C-terminal region of *P. berghei* EXP1 has been shown to interact with rodent and human Apolipoprotein H (56). While this interaction is important for *P. berghei* development within hepatocytes, it could not be recapitulated in yeast two-hybrid assays with *P. falciparum* EXP1, which displays low sequence identity with *P. berghei* EXP1 in the C-terminal region critical for interaction with Apolipoprotein H, possibly indicating lineage-specific EXP1 adaptions.

In the present study, we found that blood stage function of *P. falciparum* EXP1 is compromised by a C-terminal fusion to mNG. The topology of EXP1 orients the C-terminus on the host cytosolic face of the PVM while the N-terminus is positioned within the PV lumen (15, 16). As EXP1 bearing a bulkier mDHFR-GFP C-terminal fusion is still inserted into the PVM with proper topology (58), the EXP1-mNG defect seems unlikely to result from a perturbation in trafficking. The mNG fusion may interfere with interactions that enable EXP1 to organize into oligomeric arrays (22) as the C-terminus of *P. berghei* SEP/ETRAMP family members have been shown to be important for oligomer formation (19). However, the entire C-terminal domain of *P. berghei* EXP1 has recently been shown to be dispensable during the blood stage (although it is important in both the mosquito vector and in the hepatocyte where truncated EXP1 fails to properly traffic to the PVM, illustrating distinct EXP1 functional roles and trafficking constraints between parasite life stages) (59). The inability to similarly truncate the EXP1 N-terminus in *P. berghei* implies its essential blood stage function occurs within the PV lumen (59).

An *in silico* approach to discovery of gene function through analysis of gene relationships over large evolutionary distance suggested EXP1 may be a membrane GST (20). Subsequent *in vitro* experiments found recombinant EXP1 to possess GST activity, particularly toward hematin, leading to a model where EXP1 provides protection from the oxidative stress that results from catabolism of hemoglobin within endocytic vesicles and the digestive vacuole (20). While enzymes involved in hemoglobin degradation such as plasmepsin aspartic proteases do traffic through the PV to reach the digestive vacuole, their PV residence is transient with principal localization observed in the digestive vacuole (60). In contrast, the vast majority of EXP1 is localized at the PVM, suggesting this is the principal site of function. Here, we found that a version of EXP1 bearing an R70T mutation, which reduced recombinant EXP1 GST activity more than fivefold *in vitro,* can rescue parasite growth ∼70% as well as the wild type protein upon knockdown of endogenous EXP1 (Figure 4C). These results strongly suggest that the proposed GST activity does not fully account for the essential blood stage function of *P. falciparum* EXP1.

Known essential functions that manifest defects at the PVM include inactivation of protein export and small molecule transport following knockdown of PTEX components (8, 9, 14, 61) and RON3 (62) or a block in egress following knockdown of key players in the protease cascade that mediates PVM destruction at the end of the cycle (36, 63, 64). Parasites depleted of EXP1 do not display an export defect and develop to a late stage where they arrest mainly as trophozoites, suggesting EXP1 does not function directly in egress (Figure 5). Arrested parasites produced additional membrane structures in the host cytosol, likely reflecting alterations in the TVN, an extension of the PVM thought to function in nutrient acquisition to which EXP1 is partially localized (65–67). Parasites depleted of EXP1 also displayed an enlarged PV lumen that was often filled with hemoglobin-containing membrane-bound structures (Figure 6D). These structures are reminiscent of TVN enclosures of host cytosol (68–71) except that they are present within the PVM, and may thus reflect abnormalities in TVN formation. Alternatively, these structures might result from irregularities of the central cavity, a poorly understood structure open to the host cytosol and often observed as an indentation at the parasite periphery (72, 73). Notably, cytostomes remained visible in EXP1 knockdown parasites despite these alterations (Figure 6C,D, arrowheads). Defects in uptake of host cell cytosol would be expected to manifest in the terminal pathway compartment, the digestive vacuole, as observed with a VPS45 conditional mutant that prevents delivery of endocytosed material to the digestive vacuole, severely limiting hemoglobin degradation (74). However, digestive vacuoles appeared largely normal and hemozoin crystal formation was not grossly altered following EXP1 knockdown (Figure 5A and 6), suggesting uptake of host cell cytosol is not critically impacted.

Reduction of EXP1 levels also produced a striking change in EXP2 distribution with EXP2 often concentrated to one or two discrete points along the PVM (Figure 7). EXP2 forms a dual functional pore in the PVM that is required for small molecule transport and effector protein translocation (10, 14). Although we did not directly evaluate small molecule transport at the PVM, the fact that EXP2 pore function in protein export within PTEX is not impaired and that EXP1 knockdown parasites do not arrest at an earlier developmental stage suggests that EXP2-dependent small molecule transport is unlikely to be critically impacted.

A striking feature of the PVM is its intimate apposition to the PPM but the basis for maintaining this proximity is not known. When protein export is disabled by inactivating PTEX components, tubular distensions form and project into the host cell (14). This PV swelling appears to result from accumulation of blocked exported proteins within the PV but occurs at discrete points, with PVM-PPM anchoring still largely maintained (14). In contrast, the more uniform separation between PVM and PPM observed upon EXP1 knockdown may indicate a role for EXP1 in maintaining this intimate membrane connection, loss of which might result in broad disorganization of PVM patterning that is somehow tied to the connection with the PPM. Developmental arrest in late stage parasites may reflect a critical, though currently unclear, requirement for maintenance of this connection for proper completion of schizogony. In conclusion, our results call into question the importance of GST activity for EXP1 function and reveal a critical requirement for EXP1 in maintaining proper PVM/TVN organization.

## Acknowledgements

This work was supported by National Institutes of Health grant HL133453 to J.R.B. The funders had no role in study design, data collection and interpretation, or the decision to submit the work for publication.

We thank J. Aguiar for the EXP1 antibody, J. McBride, D. Cavanaugh and EMRR for the EXP2 antibody, C. Braun-Breton for the SBP1 antibody, W. Beatty, K. Hausmann and the WUSTL Molecular Microbiology Imaging Facility for assistance with electron microscopy, B. Vaupel for assistance with molecular cloning and E. Istvan and A. Polino for assistance with parasite culture.

## Author contributions

T.N., J.M.B., D.E.G. and J.R.B. conceived and designed the experiments. T.N., J.M.B., S.R., Y.J. and J.R.B. performed the experiments. T.N., J.M.B., S.R., Y.J., J.A.W., D.E.G. and J.R.B. analyzed the data. J.R.B. oversaw the project and wrote the manuscript. All authors discussed and edited the manuscript.

---

**Figure S1:**
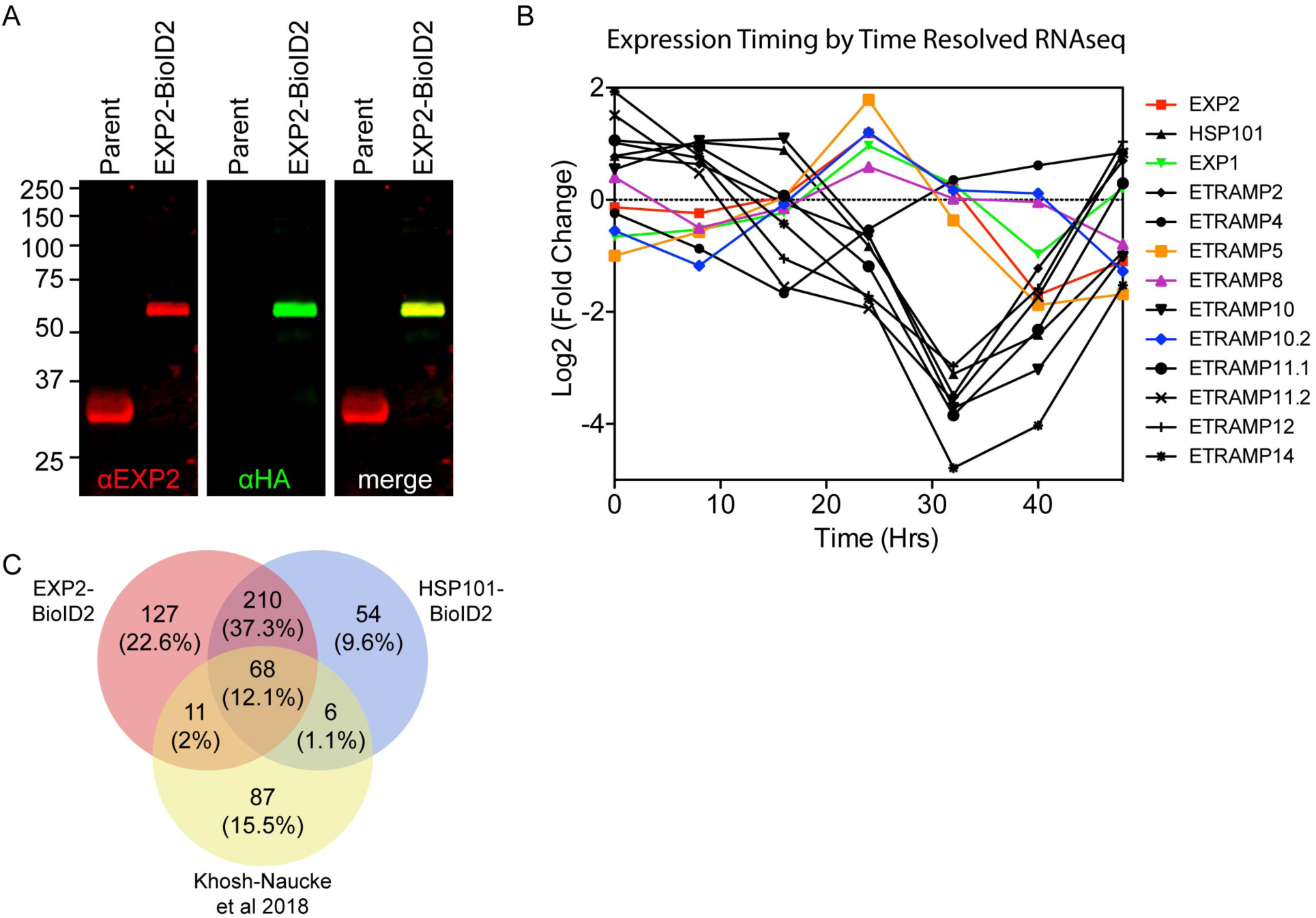
Generation of the EXP2-BioID2 fusion line and expression profiles of select hits. (A) Western blot of parent and EXP2-BioID2-3xHA parasites. Molecular weights after signal peptide cleavage are predicted to be 30.8 kDa for EXP2 and 61.2 kDa for EXP2-BioID2-3xHA. (B) Transcript fold change throughout intraerythrocytic development assessed by transcriptomic analysis of synchronized *P. falciparum* 3D7 parasites for EXP2, HSP101, EXP1 and select ETRAMP family members. Data are from RNAseq analysis by Otto and colleagues (75). Genes with similar expression pattern to *exp2* are shown in color. (C) Ven diagram summarizing overlapping and distinct proteins detected in EXP2-BioID2, HSP101-BioID2 and SP-GFP-BirA* datasets (50). Data from Khosh-Naucke et al 2018 was processed in the same way as the BioID2 datasets in this study by pooling SP-GFP-BirA* mass spectrometry datasets obtained from saponin supernatant and pellet fractions and removing all proteins identified in 3D7 negative controls.

**Figure S2:**
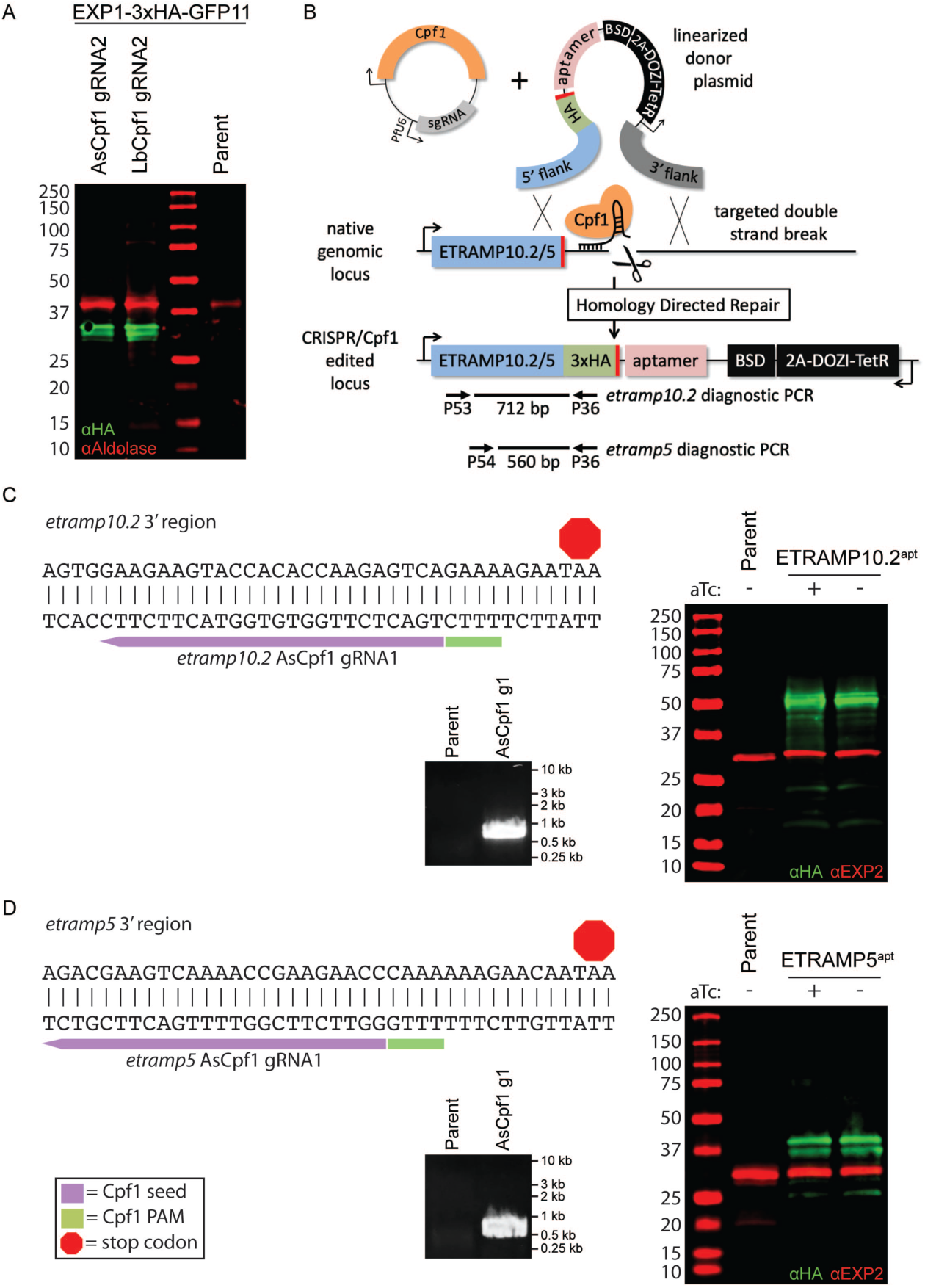
Genome editing of *exp1*, *etramp10.2* and *etramp5* with Cpf1. (A) Western blot of parent and EXP1-3xHA-GFP11 fusion parasites generated with AsCpf1 or LbCpf1 and gRNA2 as shown in Figure 2A but expressed from the selectable pUF-AsCpf1 or pUF-LbCpf1 plasmids that contain a yDHODH cassette. Aldolase serves as a loading control. Molecular weight after signal peptide cleavage is predicted to be 20.9 kDa for EXP1-3xHA-GFP11. Note that EXP1 and derivative fusions are observed to migrate at a higher molecular weight than predicted. (B) Schematic showing strategy for double homologous recombination repair of double-strand breaks mediated by Cpf1 at the 3’ end of the *etramp10.2* and *etramp5* genes to install a 3xHA fusion and 3’ TetR-DOZI-aptamers. 3’ UTR, 3’ untranslated region; yDHODH, yeast dihydroorotate dehydrogenase. (C,D) Sequence of the 3’ end of *etramp10.2* or *etramp5* with Cpf1 gRNA target indicated. Successful integration mediated by AsCpf1 to generate a 3’ fusion to 3xHA is shown by diagnostic PCR with primers indicated in the schematic and by Western blot of ETRAMP10.2^apt^ and ETRAMP5^apt^ parasites grown with or without 1µM aTc for 96 hours. EXP2 serves as a loading control. Molecular weight after signal peptide cleavage is predicted to be 39.6 kDa for ETRAMP10.2-3xHA and 19.6 kDa for ETRAMP5-3xHA. Similar to EXP1, both ETRAMP10.2-3xHA and ETRAMP5-3xHA are observed to migrate at a higher molecular weight than predicated. PAM, protospacer adjustment motif.

**Figure S3:**
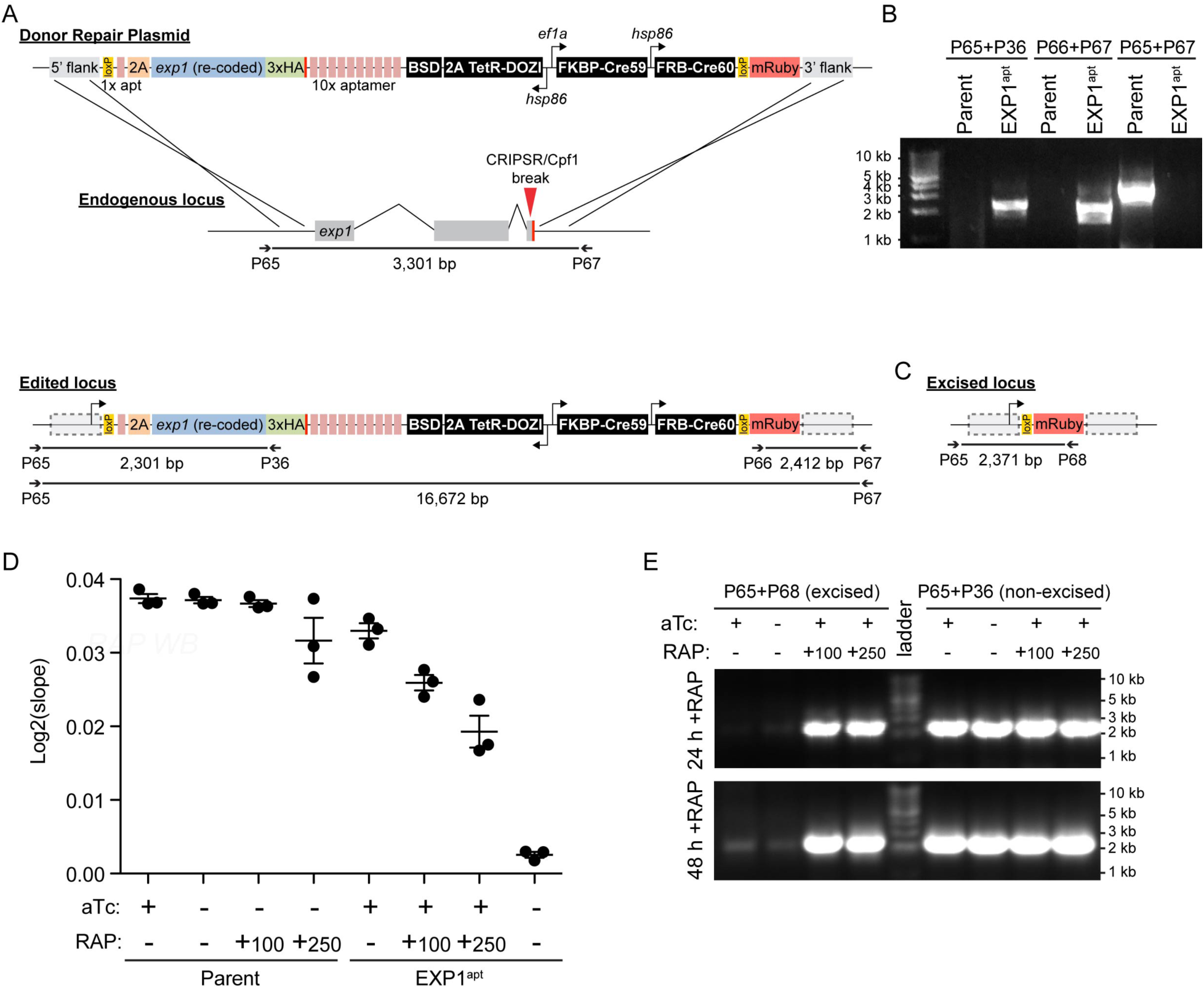
Generation of EXP1^apt^ parasites and analysis of DiCre-mediated *exp1* excision. (A) Schematic for strategy used to replace the endogenous *exp1* coding sequence with a re-coded version of *exp1* with TDA and DiCre elements by Cpf1 editing and double homologous recombination. (B) Diagnostic PCR with primers indicated in the schematic in (A) showing successful integration at the 5’ and 3’ ends of the *exp1* locus in EXP1^apt^ parasites. The absence of the product in EXP1^apt^ using primers P65/P67 is likely due to the very large amplicon size. (C) Schematic of *exp1* locus following excision between *loxP* sites by DiCre. The pEXP1^apt^ plasmid was designed so that DiCre excision of the modified locus would place the promoter-less *mruby3* coding sequence under the control of the endogenous *exp1* promoter. (D) Growth analysis of parental and EXP1^apt^ parasites with or without 1 µM aTc or with and without 100 µM or 250 µM rapamycin. The scatter plot shows the slope of the line fitted to the mean of log2-transformed parasitemias for each of three independent experiments. Error bars indicate SEM. (E) Diagnostic PCR for *exp1* excision by DiCre with primers indicated in the schematics.

